# Optical flow analysis reveals that Kinesin-mediated advection impacts on the orientation of microtubules in the *Drosophila* oocyte

**DOI:** 10.1101/556043

**Authors:** Maik Drechsler, Lukas F. Lang, Layla Al-Khatib, Hendrik Dirks, Martin Burger, Carola-Bibiane Schönlieb, Isabel M. Palacios

## Abstract

The orientation of microtubule networks is exploited by motors to deliver cargoes to specific intracellular destinations, and is thus essential for cell polarity and function. Reconstituted *in vitro* systems have largely contributed to understanding the molecular framework regulating the behavior of microtubule filaments. In cells however, microtubules are exposed to various biomechanical forces that might impact on their orientation, but little is known about it. Oocytes, which display forceful cytoplasmic streaming, are excellent model systems to study the impact of motion forces on cytoskeletons *in vivo*. Here we implement variational optical flow analysis as a new approach to analyze the polarity of microtubules in the *Drosophila* oocyte, a cell that displays distinct Kinesin-dependent streaming. After validating the method as robust for describing microtubule orientation from confocal movies, we find that increasing the speed of flows results in aberrant plus end growth direction. Furthermore, we find that in oocytes where Kinesin is unable to induce cytoplasmic streaming, the growth direction of microtubule plus ends is also altered. These findings lead us to propose that cytoplasmic streaming - and thus motion by advection – contributes to the correct orientation of MTs *in vivo*. Finally, we propose a possible mechanism for a specialised cytoplasmic actin network (the actin mesh) to act as a regulator of flow speeds; to counteract the recruitment of Kinesin to microtubules.

**HIGHLIGHT SUMMARY:** Cytoskeletal networks do not exist in isolation, but experience crowded and dynamic intracellular environments. However, microtubule-environment interactions are not well understood, and such system-environment interactions are an unresolved question in biology that demands bridging across disciplines. Here we introduce an optical flow motion estimation approach to study microtubule orientation in the *Drosophila* oocyte, a cell displaying substantial cytoplasmic streaming. We show that microtubule polarity is affected by the regime of these flows, and furthermore, that the presence of flows is necessary for MTs to adopt their proper polarity. With these findings we are contributing to further understanding how microtubules organize in their impacting natural environment.

## INTRODUCTION

Eukaryotic life depends on many dynamic processes, including for example cell division, cell migration, and cell polarization. These processes in turn strongly rely on highly organized microtubule (MT) arrays. All MT networks are polarized, with the minus end of each filament linked to a nucleating centre (MT organising centre or MTOC), and the plus end growing away from these centres. This polarity is exploited by specific motor proteins to transport cargoes along MTs in a defined direction, and is thus essential for the function of MT networks.

A number of biophysical studies, using reconstituted *in vitro* systems, have helped to understand the mechanical properties of MTs, setting the stage for deciphering the behavior of MTs *in vivo*. However, much needs to be learnt about the properties of MTs in their natural intracellular environment. For example, a rather new concept emanating from *in vivo* experiments is that controlling nucleation and the position of minus ends of single filaments alone is not always sufficient to establish the proper polarity of the entire MT network. MT plus ends must be controlled as well in order to allow motor proteins to deliver their cargoes to the correct destination. Plus ends can be regulated at various levels, including dynamic instability, capturing, and direction of growth. Dynamic instability describes a process in which MT polymerisation is interrupted by a rapid depolymerisation phase, followed by a ‘rescue’ process(Mitchison and Kirschner, 1984). Various MT-associated proteins, such as motors and MT plus end-tracking proteins (+TIPs), are known to regulate dynamic instability (Akhmanova and Steinmetz, 2015). MT plus ends can also be captured at the cell cortex, a process also involving +TIPs and motors, such as the Dynein/Dynactin complex (Nieuwburg *et al*., 2017) (and reviewed in (Akhmanova and Steinmetz, 2015)). However, very little is known about how the direction of growth of plus ends, and therefore the orientation of MTs, is controlled in cells. In axons, adenomatous polyposis coli (APC) seems to regulate plus end direction (Purro *et al*., 2008), while Fidgetin-like1, a MT-associated ATPase, controls both dynamics and plus ends direction (Fassier *et al*., 2018). MT bending also impacts on the direction of plus tip growth, as the MT tip rotates due to local bend formation (Kent *et al*., 2016). Furthermore, +TIPs that contain actin-binding motifs can influence MT growth direction by guiding dynamic plus ends along actin bundles (Kodama *et al*., 2003; Jiang *et al*., 2012).

In the mid-oogenesis stage 9 (st9) oocyte of *Drosophila melanogaster*, MTs are nucleated from the antero-lateral cortex in a gradient of diminishing abundance toward the posterior pole, where nucleation is absent. In addition, the growing plus ends exhibit a weak posterior orientation bias (Doerflinger *et al*., 2006; Parton *et al*., 2011; Nashchekin *et al*., 2016). This weak polarization bias of plus ends towards the posterior pole of the oocyte is sufficient and necessary for the localization of body plan determinants to the very ‘posterior tip’, as well as the formation of the pole plasm (needed for germ cell development) in this region. For example, the plus end motor Kinesin-1 (Kin from here on) localizes *oskar* mRNA to the ‘posterior tip’ of the st9 oocyte, an essential step in the establishment of the anterior-posterior (A-P) axis and the formation of the pole plasm (Brendza *et al*., 2000).

In addition to a polarized MT cytoskeleton, the oocyte displays a Kin-dependent bulk motion of the cytoplasm called cytoplasmic streaming or flows (Palacios and St Johnston, 2002; Serbus *et al*., 2005; Ganguly *et al*., 2012; Drechsler *et al*., 2017). Compared to st9, late stage oocytes (st11) exhibit faster and larger scale flows that are induced not only by Kin-mediated viscous drag of transported cargo (similar to st9), but also by Kin-mediated MT-sliding (Lu *et al*., 2016; Monteith *et al*., 2016). At these later developmental stages, these faster flows are important for mixing the cytoplasm of the large oocyte (Ganguly *et al*., 2012), and aiding the asymmetric localization of developmental determinants and mitochondria (Forrest and Gavis, 2003; Hurd *et al*., 2016; Lu *et al*., 2018). In addition, mathematical modeling has suggested that faster flows can induce parallel MT arrays when the MTs are near a barrier (such as the cortex)(Monteith *et al*., 2016). However, in st9 oocytes, when posterior cargoes such as *oskar* mRNA are first localized, MT sliding has been excluded as the source of cytoplasmic streaming (Lu *et al*., 2016). Cytoplasmic flows at st9 are slowed down by a cytoplasmic actin mesh. When this actin mesh dissolves during mid-oogenesis, the flows become faster and more organized, and the MT arrange into sub-cortical bundles (Dahlgaard *et al*., 2007; Quinlan, 2013). However, the mechanism by which the actin mesh regulates flows is unknown. Furthermore, it is unclear whether cytoplasmic flows at st9 have an impact on the organization of the MT cytoskeleton. Recently, we found that st9 flows constitute a key force driving the persistent motion of vesicles and actin filaments (Drechsler *et al*., 2017). These observations prompted us to address the question of how advection (active transport induced by fluid flows) impacts on the polarization of the MT network in st9 oocytes. In this way, we aim to contribute to the insufficient knowledge on how the direction of MT growth is controlled *in vivo*.

In order to assess the global MT orientation in st9 oocytes, and to investigate the growth direction of MT plus ends *in vivo*, we used EB1::GFP (EB1 from here on). EB1 exclusively decorates the growing plus end of MTs, resulting in dynamic ‘comets’ moving through the cytoplasm at a speed of ~100-600 nm/sec across various cell types, including the oocyte (at an average and maximum speeds of 230 nm/sec and 600 nm/s, respectively (Parton *et al*., 2011; Nieuwburg *et al*., 2017). Analyzing the orientation of MTs in complex networks has proven technically challenging, and requires suitable imaging and image analysis tools. Especially for the *Drosophila* oocyte, we found the published method too demanding on the imaging level, requiring state of the art wide-field deconvolution microscopy and elaborate image processing (Parton *et al*., 2011), and thus unfeasible for the various experimental conditions that our study required. Here, we developed an image analysis strategy that allows an efficient characterisation of direction and distribution of EB1 comets from confocal image series by an optical flow (OF)-based motion estimation approach. In general, OF allows to estimate the apparent motion of objects or other intensity variations in a sequence of images (Horn and Schunck, 1981). In addition, variational OF methods constitute a well-established framework for dense motion estimation, and do not require elaborate segmentation or tracking of the studied structures. OF methods outperform popular methods, such as particle image velocimetry (PIV), for motion analysis in certain settings and, in particular, in the presence of noise (Ruhnau et al., 2005; Vig *et al*., 2016). While variational OF methods have been used predominantly to investigate the dynamics of entire cells (Amat *et al*., 2013; Boric *et al*., 2013; Guo, 2014), recent works focused also on intracellular motility (Delpiano *et al*., 2012; Frerking *et al*., 2014; Vig *et al*., 2016; Boquet-Pujadas *et al*., 2017; Huang *et al*., 2017).

In this study, we investigated how cytoplasmic streaming influences the polarization of the MT cytoskeleton. For this, we introduce a two-step image analysis approach that is based on image denoising and variational OF, and is able to estimate approximate velocities (speed and direction) of EB1 motion in confocal image sequences. This approach will assist many researchers interested in characterizing MT polarity from confocal images of their tissue of choice. Our findings revealed that cytoplasmic streaming is necessary and sufficient to regulate the polarity of the MT network in st9 oocytes. Furthermore, our data suggests that the actin mesh regulates the recruitment of Kin to MTs, indicating a new mechanism by which the actin cytoskeleton influences MT-based transport and advection. With this work, we further contribute to understanding the *in vivo* properties of MTs, and the interactions of MTs with their natural environment.

## RESULTS

### Quantification of MT plus-tip directionality by optical flow analysis

To study the spatial orientation of MT filaments *in vivo*, we used oocytes expressing EB1 (Figures 1, 3, 4 and 5). EB1 constitutes a marker for growing MT plus-tips and has been used in fly oocytes before (Parton *et al*., 2011; Nieuwburg *et al*., 2017). In these previous studies, wide-field deconvolution microscopy was used to image EB1 dynamics in various areas of st9 oocytes. However, wide-field microscopy has a limited focus depth, only allowing to image EB1 dynamics close to the cortex of the relatively large oocyte, and requires a complex acquisition procedure. In order to improve imaging depth and simplify the acquisition procedure, we implemented a strategy that combines conventional confocal microscopy with image denoising and motion analysis by variational OF analysis.

**Figure 1.**
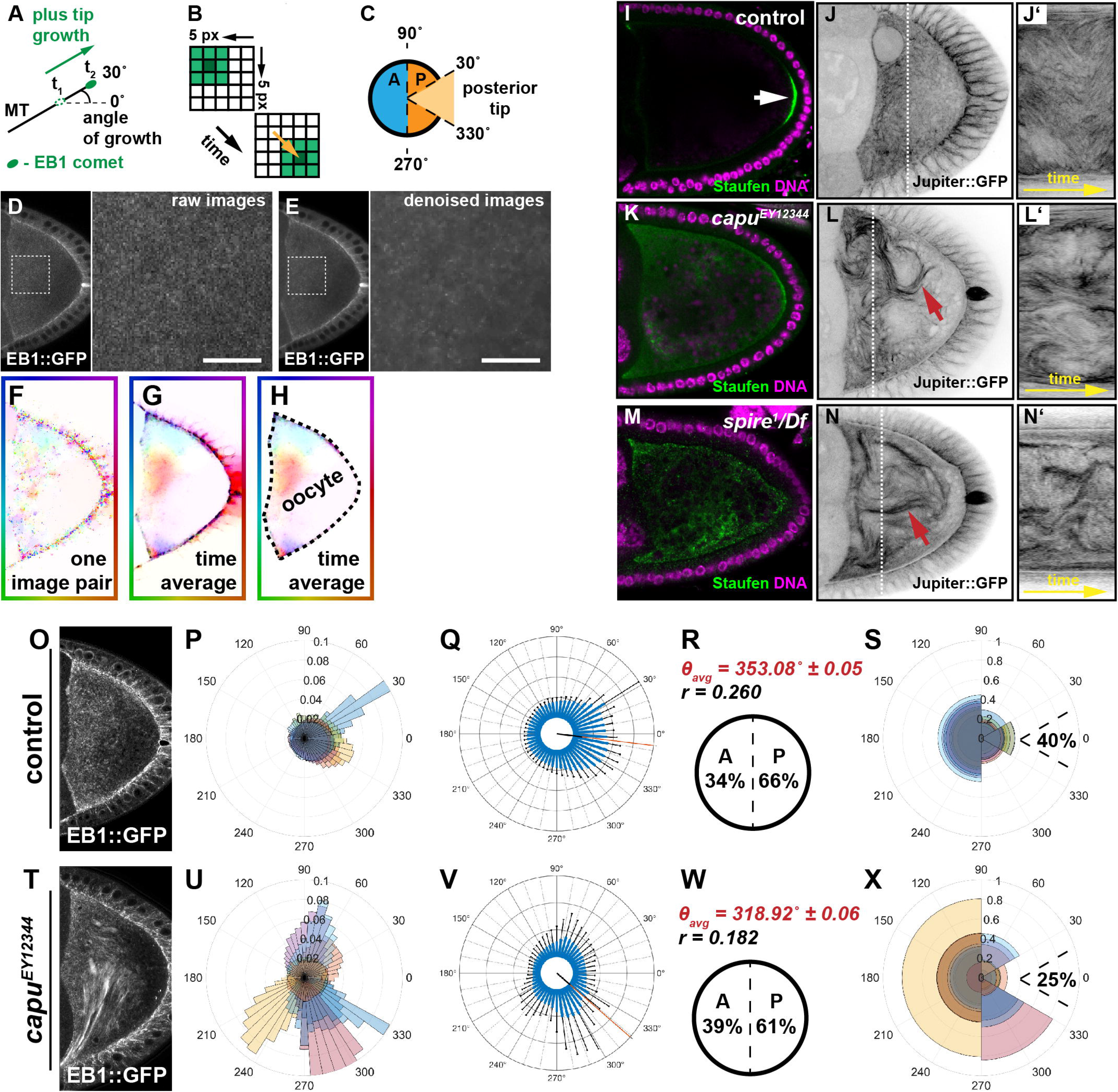
MT orientation is aberrant in oocytes with faster flows. **A)** Schematic representation of MT orientation. EB1 specifically associates with the growing end (plus-end) of MTs and therefore serves as read-out of their spatial orientation. The orientation of MT growth is represented as an angle, deviating from an imaginary anterior (180°) to posterior axis (0°). **B)** Simplified scheme, illustrating the underlying principle of OF-based motion estimation. Shown are two consecutive frames of a 5 x 5 pixel wide image sequence that contains a rectangular object of different pixel intensities - from light green (little signal) to dark green (maximum signal). OF assumes that the intensities of a signal do not change along its trajectory. Based on this assumption, variational OF allows to estimate a displacement vector for each pixel (the yellow arrow shows the displacement vector of the centre pixel of the object). **C)** Definition of growth directions based on OF-estimated velocities. Angles of velocities between 90°-270° are regarded as anterior (A, blue hemicircle), the complementary set of angles as posterior (P, orange hemicircle). The sub-population of all posterior signals (orange) that fall between 330°-30° (light orange sector) are considered to grow towards the ‘posterior tip’. **D)** Single frame of an unprocessed image sequence (raw data) showing an oocyte expressing EB1. The magnified area is indicated by a dashed box. **E)** Same frame as shown in (D), after applying the denoising step (Movie S2). Scale bars are 10 μm. **F)** Shown is the OF (displacement vector) field between two frames of the image sequence in D. **G)** Average OF (over the entire sequence). **H)** Hand-drawn segmentation mask of the oocyte. For the analysis, only the displacement vectors within this segmentation were considered. **I)** Staufen (green) localization in control oocytes (arrow). The protein localizes in a tight posterior crescent by the end of st9. **J)** Living control oocyte expressing the MT-binding protein Jup (Movie S5). **J’)** Kymograph/space-time plot (along the indicated dotted line in C), showing dynamic bending of MTs over time. **K and M)** Staufen fails to localize in *capu* or *spire* mutants and distributes throughout the cytoplasm of the cell. **L and N)** Living *capu* (L) or *spire* (N) mutant oocytes, expressing Jup. Fast cytoplasmic flows lead to the formation of dense and long MT bundles (red arrows), as well as increased MT bending (Movie S5). **L’ and N’)** Kymograph (along dashed line in L and N) indicate a stronger displacement of MT bundles over time in *capu* (L’) or *spire* (N’) mutants. **O)** Standard deviation (temporal) projection of EB1 comets in a control image sequence (in total 65 s). **P)** Rose diagram (angular histogram) with 50 bins depicting the distribution of EB1 directions in individual control cells within the corresponding segmented oocyte. Each colour represents the angular histogram of the directions from one oocyte. **Q)** Same data as shown in (P) with angular histograms averaged over all cells (n=8). Error bars (in black) indicate the standard deviation for each bin (in blue). **R)** Mean angular direction θ_avg_ of the histogram shown in (Q) (also indicated by a red line in (Q)) and the length *r* (between 0 and 1) of the mean resultant vector (length of black line in (Q) originating from the centre), which relates to the circular variance *S* = 1 – *r* of the distribution shown in (Q). Anterior-posterior bias of all EB1 growth directions. **S**) Rose diagram similar to (P) for control cells but with growth directions binned into four bins (30°-90°, 90°-270°, 270°-330°, and 330°-30°). Moreover, the fraction of posterior-growing EB1 comets pointing towards the ‘posterior tip’ (330°-30°) is indicated. **T)** Standard deviation (temporal) projection of EB1 comets in an image sequence of a *capu* mutant oocyte (in total 65 s). **F-I)** Quantified EB1 directions in *capu* cells, following the same experimental pipeline described for the control cell.

**Figure 2.**
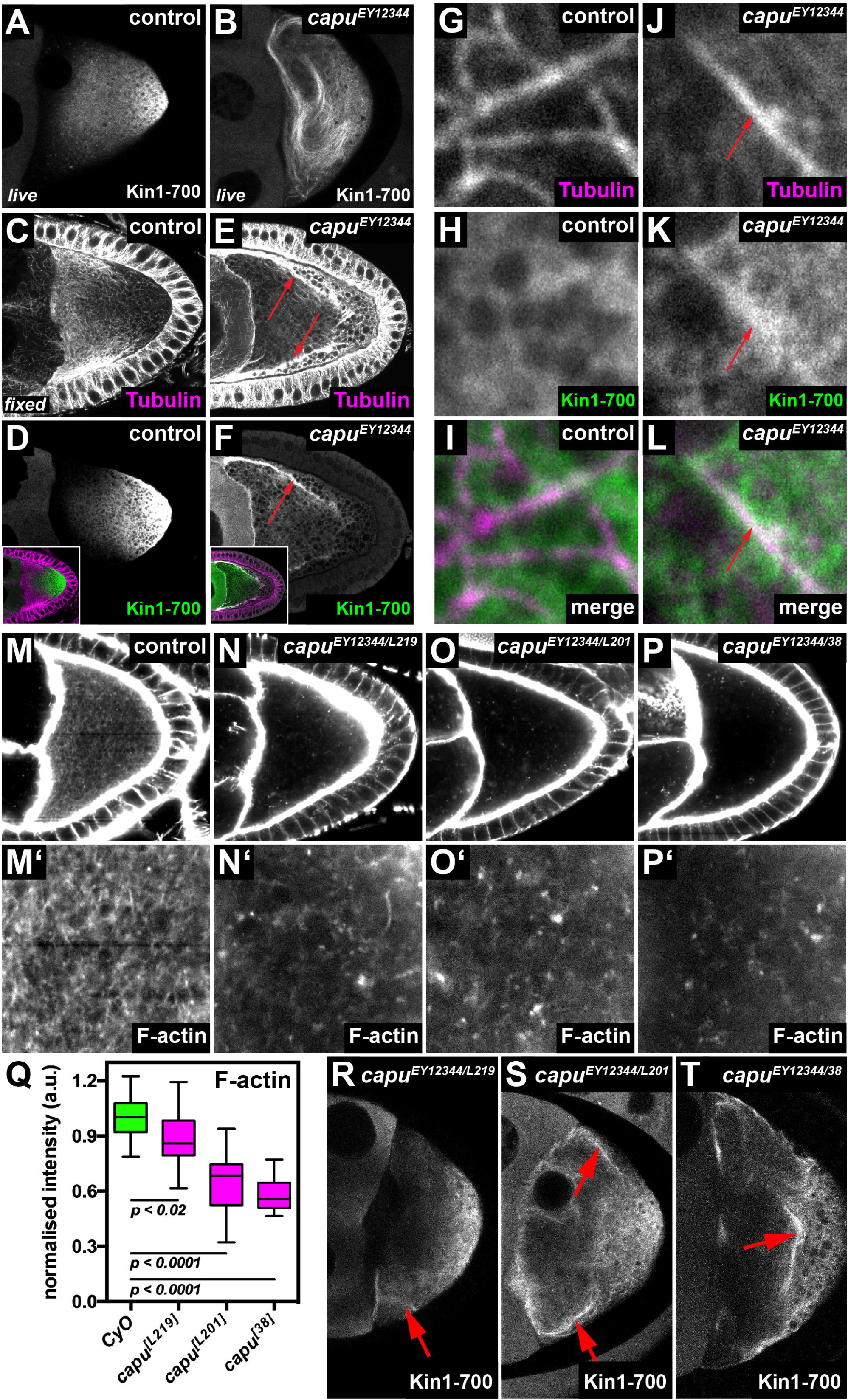
The cytoplasmic actin mesh counteracts the recruitment of Kinesin to MTs. **A,B)** Living control (*capu^EY12344^/CyO* in A) and *capu* mutant oocyte (*capu^EY12344^/capu^EY12344^* in B), expressing Khc::GFP. While the fusion protein mainly localizes posteriorly in control cells (A), it strongly decorates MTs in *capu* mutant cells (B and Movie S10). **C-F)** Fixed control (*capu^EY12344^/CyO*) oocytes (C,D) and *capu* mutant cells (*capu^EY12344^/capu^EY12344^*)(E,F), expressing Khc::GFP. Cells were stained against αTubulin (magenta) and the Kin1-700-GFP fusion protein (green). In controls, Tubulin and Khc::GFP localize to opposed gradients and show little overlap (C,D). Conversely, in *capu* mutants, Khc::GFP strongly co-localizes with Tubulin (red arrows in E,F). Insets in D and F show merged channels. **G-L)** Higher resolution images of fixed control (*capu^EY12344^/CyO*) (G-I) and *capu* mutant (J-L) cells, stained against αTubulin (magenta) and GFP (green). While Khc::GFP localizes diffusely around MTs in control cells (G-I), it strongly co-localizes to MTs in *capu* mutants (red arrows in J-L). **M-P)** The cytoplasmic actin mesh in control (M, *capu^EY12344^/CyO*) and transheterozygous *capu^EY12344^/capu^L219^* (N), *capu^EY12344^/capu^L201^* (O), and *capu^EY12344^/capu^38^* (P). **Q)** Quantification of signal intesities of the actin mesh as shown in (M-P). All cells harbor the strong hypomorphic allele *capu^EY12344^* over either CyO (n=15), *capu^L219^* (n=21), *capu^L201^* (n=20) or *capu^38^* (n=21). **R-T)** Khc::GFP expressed in living *capu^EY12344^/capu^L219^* (R), *capu^EY12344^/capu^L201^* (S), and *capu^EY12344^/capu^38^* (T) oocytes. Red arrows indicate Khc::GFP positive MT bundles.

**Figure 3.**
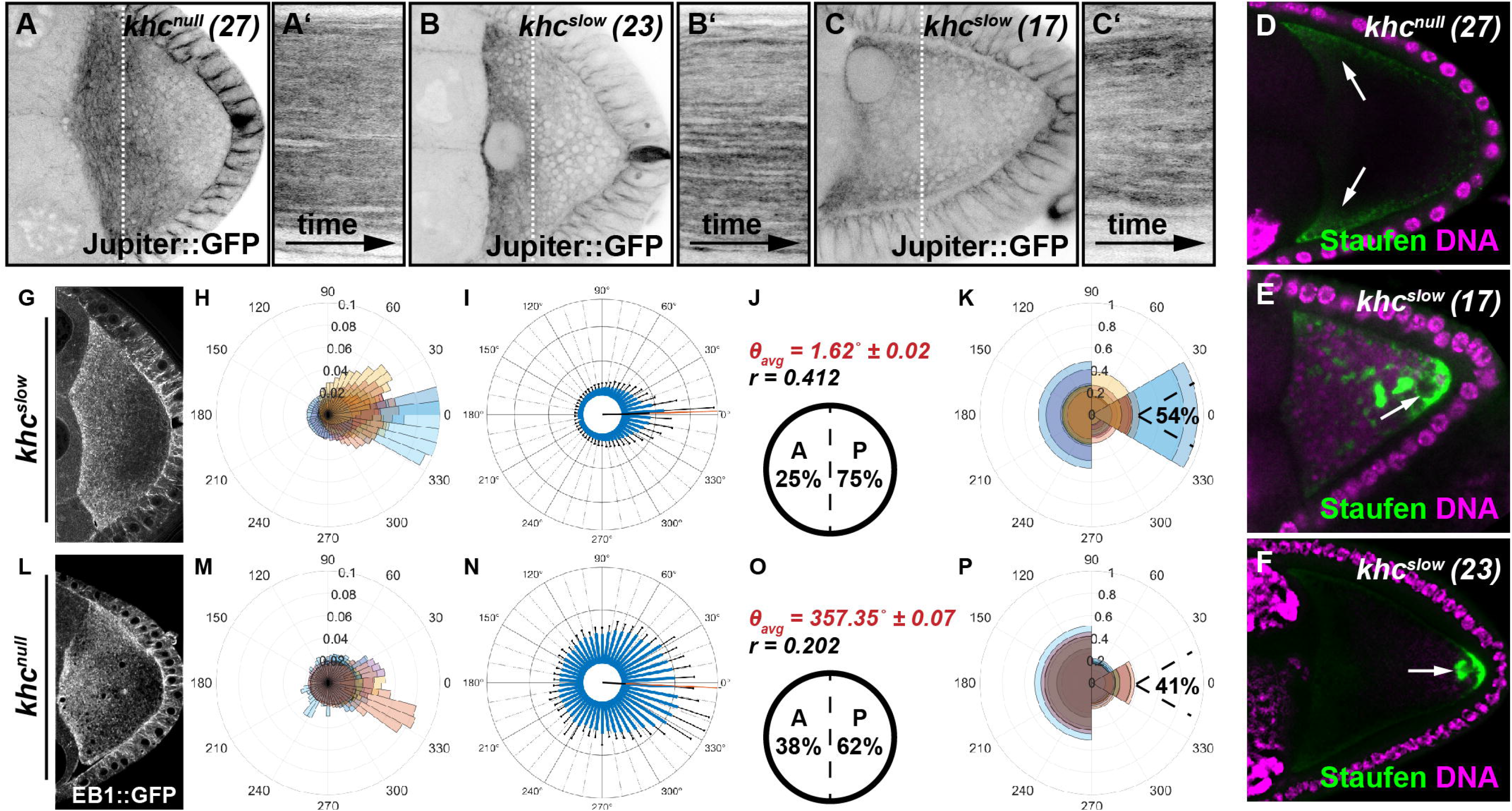
Kin activity impacts on the spatial orientation of MTs in the oocyte. **A-C)** Still frames and kymographs of live oocytes expressing Jup. Cells harbor a null mutation (*khc^null^ (27)*), or single point mutations in the motor domain (*khc^slow^ (23)* and *khc^slow^ (17)*), rendering the motor slower. Compared to controls, all *khc* mutant cells exhibit no cytoplasmic flows and thus no MT bulk motion (Movie S6). **D-F)** Posterior cargo localization in different *khc* mutant alleles. Staufen (green) is not transported to the posterior in cells lacking Kin (*khc^null^ (27)*) and is found in the anterior corners of the cell (arrows). In contrast, in both of the slow Kin alleles (*khc_null_ (23)* and *khc^slow^ (17)*) a considerable amount of Staufen becomes transported to the posterior. However, compared to controls (Figure 2A) Staufen is not only localized in a posterior crescent, but also in dots within the posterior cytoplasm (arrows). **G-P)** OF analysis of EB1 growth directionality in *khc^slow^* (G-K, n=10) and *khc_null_* (L-P, n=10) oocytes, following the same experimental pipeline described in Figure 1.

**Figure 4.**
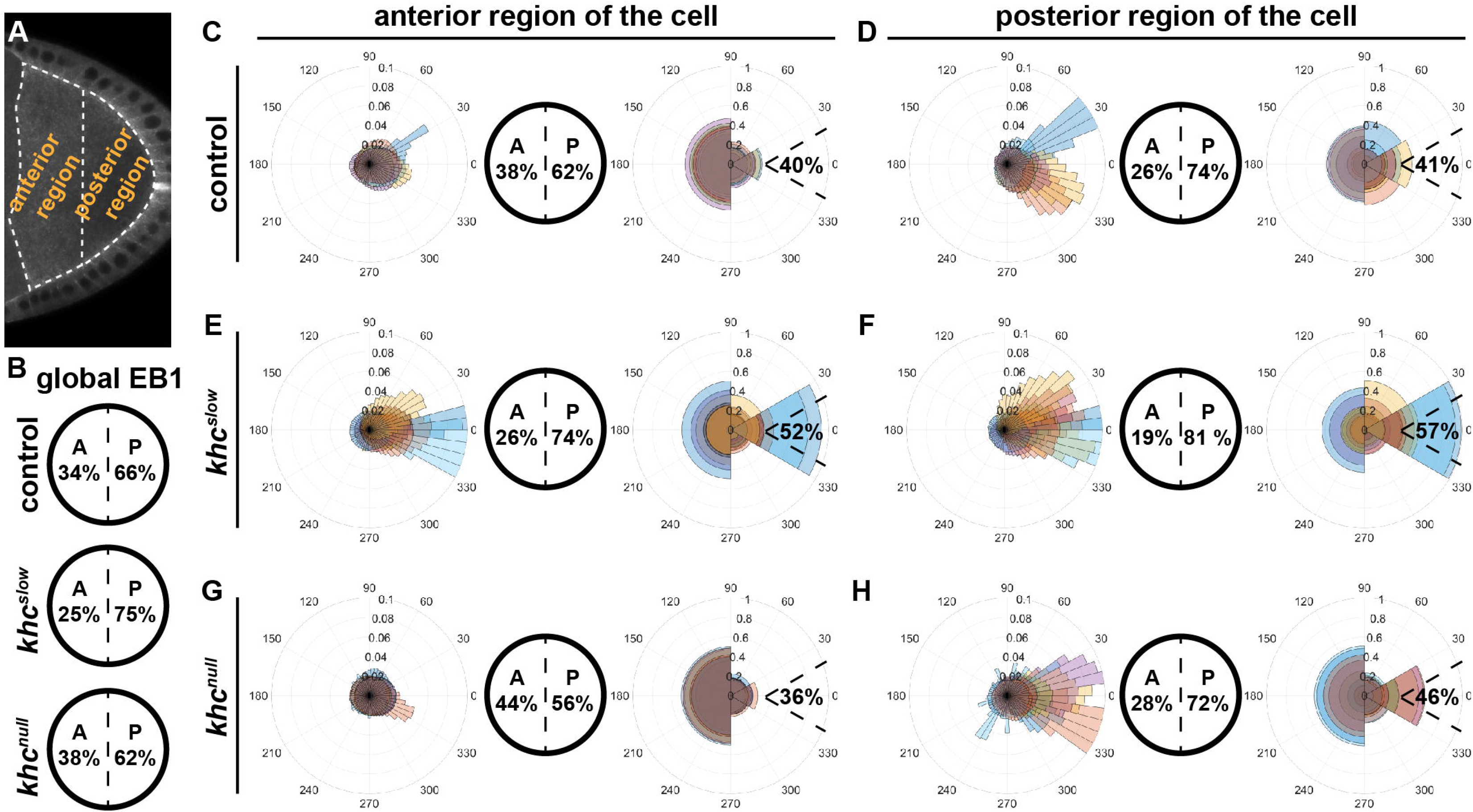
Anterior and posterior regional impact of Kin activity on MT orientation. **A)** Single frame of an oocyte expressing EB1. Dashed lines represent the anterior and posterior regions of the oocyte used to extract orientation data. **B)** Global EB1 signal directions within the entire oocyte (as shown in Figure 1 and 3). **C-H)** In each case, from left to right: distribution of EB1 growth orientation visualized in a rose diagram, the anterior-posterior orientation, and the ‘posterior tip’ orientation. **C,D)** Data for control cells (n=8). **E,F)** Data for *khc^slow^* mutant cells (n=10). **G,H)** Data for *khc^null^* mutant cells (n=10).

OF-based motion estimation relies on the assumption that particles approximately maintain their pixel intensity as they move. In thick biological samples the intensity of fluorescent signals declines with the depth of the imaging plane. To minimize the effects of tissue thickness on signal intensity, we imaged within a single focal plane in the middle of the oocyte over the course of at least one minute (for details see Material and Methods). With this approach, we avoided measurable bleaching and ensured that EB1 comets would only be detected within a thin focal plane of ~1.3 μm. Thus, comets moving orthogonally to the focal plane would be lost instead of getting dimmer or brighter. As a consequence, our analyzes capture the 2D motion of EB1 comets within a 3D intracellular environment. However, the orientation of individual oocytes on the microscope slide is random. Together with averaging across several cells, this results in 2D data obtained from various angles (one precise orientation per cell), which indirectly allows to acquire information about the 3D behavior. As OF is formulated on a per-pixel basis, this method allows inferring a displacement vector for each pixel and does not require sophisticated tracking of individual particles (Figure 1A and B). Since MTs grow one order of magnitude faster compared to the speed of cytoplasmic flows in st9 oocytes, we took one image every 650 ms, resulting in a loss of spatial resolution, and causing the raw data to exhibit considerable high noise (Figure 1D). Due to this high noise level, preceding denoising of the image data was required prior to applying the motion estimation. We found that total variation-based (Rudin et al., 1992) image denoising with additional temporal regularization was sufficient to improve the signal-to-noise ratio and the visibility of EB1 comets (Figure 1E, Figure S1-S3 and Movie S1).

In the next step, displacement vector fields were computed with a variational OF method (Figure S4). After accounting for the pixel size and the time interval between consecutive frames, these displacements can be regarded as approximate velocities of EB1 motion (Figure 1F-H). Importantly, the acquired velocities are a sum of all motion experienced by the imaged EB1 signal, including filament growth and MT displacement by advection and transport. A typical image sequence of 100 frames and a 512 x 256 pixel resolution results in a total number of approximately 13 million computed vectors that require appropriate interpretation. The computation time for processing one typical sequence amounted to less than 25 minutes on average.

We visualized the estimated vector fields with the help of a standard colour-coding (Baker, 2011). The vector at each pixel and at a certain frame is represented by a colour that is determined by the direction of the signal’s movement (see the colour-coding at the boundary of the images in Figure 1F-H). The colour’s intensity is determined by the relative velocity of the signal. The stronger the intensity, the higher the velocity. For our statistical analyzes, we considered only vectors located within a hand-drawn segmentation mask that outlines the oocyte in each sequence (Figure 1H). The direction of each vector could then be represented in polar coordinates. The distribution of the angles obtained from each individual cell is visualized in a rose diagram (Figure 1O and P). In addition, we show the histogram of the distributions averaged over all cells of a given genotype (Figure 1Q).

We then used OF to describe the MT orientation in control oocytes (Figure 1O-S, Figure S5). Since all cells were orientated with the posterior pole to the right during imaging, the angles given in the rose diagram reflect the directional movement of EB1 comets. For a quantitative description of MT orientation, we determined the mean angular direction of the estimated motion of EB1 signals (*θ*_avg_, considering all recorded cells of one genotype, Figure 1R, Table 1) and the frequency of movements directed towards the anterior (all within the region 270° to 90°, blue hemicircle in Figure 1C) or the posterior (all within the region 90° to 270°, orange hemicircle in Figure 1C). As a descriptor of how focused MTs grow towards the very ‘posterior tip’, we also determined the relative frequency of posterior-directed EB1 signals that fall within a sector of 60° (from 30° to 330°), called ‘posterior tip’, inspired by the region where determinants localize and pole plasm forms (light ogange sector in Figure 1C). This ‘posterior tip’ frequency only considers EB1 signals that were previoulsy found to grow towards the posterior (orange hemicircle in Figure 1C), and thus the ‘posterior tip’ value represents a sub-population of all posterior signals that grow with an angle between 30° to 330°. In summary, we find that all growing MTs exhibit a global (over the entire oocyte) posterior orientation bias, with 66% of all comets growing towards the posterior of control cells (Figure 1R). These findings are in agreement with previous reports of biased MT orientation in st9 oocytes (Parton et al., 2011), and provide a first level of validation to establish OF as a reliable way to investigate the orientation of MTs. In addition, we find that 40% of the 66% of comets that grow towards the posterior are oriented towards the ‘posterior tip’, where determinants localize and the pole plasm forms (Figure 1S).

**Table 1.**
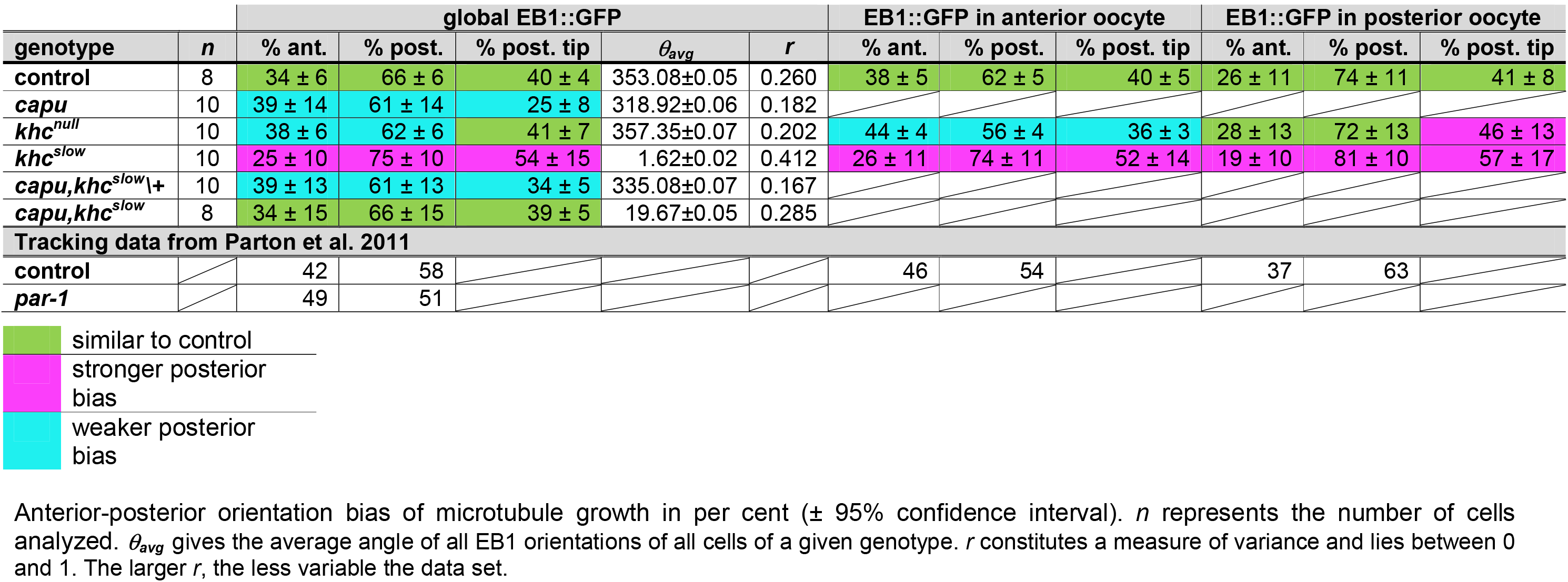
Summary of EB1 orientation data, extracted from confocal time series by variational OF analysis.

### OF is sufficient to detect changes in microtuble network orientation

In order to test whether OF-based approaches are able to capture EB1 directionality in other cell types, we investigated EB1 comets in the follicular epithelium surrounding the egg chamber. In these epithelial cells, MTs are nucleated apically and grow predominantly towards the basal membrane, parallel to the lateral membranes of the cell. Notably, single comets can easily be identified and followed by eye, which allowed us to assess the accuracy of the OF results. OF analysis captures the directionality of MTs in these follicle cells accurately, supporting the suitability of the approach to estimate MT orientation *in vivo* (Movie S2 and Figure S6-S7).

We next tested whether OF can pick up changes in the architecture of MT neworks in the oocyte, and analyzed cells harboring a mutation in *gurken* (*grk*), which exhibit MT organization and cell polarity defects (Gonzalez-Reyes *et al*., 1995; Roth *et al*., 1995; Januschke *et al*., 2002). With our OF analysis we were able to detect an aberrant MT organization in *grk* oocytes, with EB1 comets showing a variable orientation from cell-to-cell (Movie S3 and Figure S6). Although, the number of tested cells is low, and thus the actual biological meaning of these findings needs further investigation, our results confirm that OF is able to detect aberrations in MT networks in complex systems like the oocyte.

### Aberrantly fast cytoplasmic flows change MT motion, bundling and orientation

In order to investigate the relationship between cytoplasmic advection and the organization of the MT network, we monitored the bulk behavior of MTs in control cells and in oocytes with faster flows. Oocytes mutant for *cappuccino* (*capu*) or *spire* display faster flows at st9 (Figure S9A, B) and fail to localize posterior cargoes like the mRNA-binding protein Staufen (Figure 1I, K and M) (Theurkauf, 1994; Dahlgaard *et al*., 2007; Quinlan, 2013). Capu (a formin) and Spire are actin nucleators that are required for the formation of a cytoplasmic actin mesh that regulates the magnitude of flows by a yet unknown mechanism. It has been shown in fixed samples that the removal of the actin mesh seems to cause bundeling of MTs in close proximity to the cell cortex. However, dynamic MT behavior in living mutant oocytes has not been addressed in detail.

To monitor the bulk movement of MTs, we imaged cells expressing the MT associated protein Jupiter::GFP (Jup), which labels the entire MT filament. Compared to control cells, we observed that MTs in homozygous *capu^EY12344^* or transheterozygous *spire^1^/Df(2L)Exel6046* mutant oocytes appear as thick bundles, which dramatically bend and buckle (Figure 1I-N’ and Movie S5). Such higher ‘packing’ of filaments does not necessarily indicate their crosslinking, but argues that the high hydrostatic pressure in fast flowing oocytes is sufficient to bundle MT filaments. More importantly, and in contrast to previous descriptions, our data demonstrate that in *capu* and *spire* oocytes MTs cluster even when they are not in close proximity to the cortex, and thus that bundles can form even when the MTs are not pushed against the cortex. Furthermore, the observed MT bundles appear longer when compared to control cells, extending further into the posterior (red arrows in Figure 1L and N). In summary, these results indicate that changing the regime of cytoplasmic advection impacts on the bulk motion of MTs, as well as on their cytoplasmic bundeling state.

In *capu* and *spire* mutant oocytes flows can reach speeds up to 200 nm/s, well within the regime of plus tip growth. In such mutant situation, the plus tip of single MTs might not be able to ‘outgrow’ the effect of cytoplasmic advection, and thus flows might become a dominant contributor of MT orientation. To test whether faster flows alter the orientation of MTs, we quantified the directionality of EB1 signals in *capu* mutant cells with our OF approach, and found substantial alterations (Figure 1T-X). Compared to controls, *capu* mutants exhibit large amounts of comets in close proximity to each other, which move into the same direction. This suggets that *capu* mutant oocytes display arrays of parallel growing filaments (Figure 1T and Movie S4). The motion of such EB1 ‘arrays’ (which varies from cell to cell, Figure 1T-X), is likely to correlate with the bundling of MT filaments described above, and could be interpreted as a direct result of faster flows. Interestingly, the global posterior EB1 orientation bias was only mildly decreased in most *capu* mutant oocytes (Figure 1V, W vs. Figure Q, R and Table 1). However, MTs are less focused towards the posterior tip in *capu* mutants, with only 25% of the posterior-directed EB1 signals growing with an angle between 30° to 330°. These findings are in a good agreement with the determinant mis-localization defects in *capu* and *spire* mutant cells (Figure 1X vs. Figure 1K and M).

In summary, these findings together demonstrate that the lack of the actin mesh, and the resulting increase in flows speeds, impact on the orientation of MTs, and lead us to speculate that at mid-oogenesis flows must be kept in a precise regime for molecular determinants to be localized correctly. However, how the actin mesh actually regulates the magnitude of flows in the oocyte is not known

### The cytoplasmic actin mesh counteracts the recruitment of Kinesin to MTs *in vivo*

The MT plus-tip directed motor protein Kin constitutes the major driving force of cytoplasmic flows in the *Drosophila* oocyte (Palacios and St Johnston, 2002). To start investigating possible mechanisms by which the actin mesh regulates the magnitude of cytoplasmic flows, we monitored the distribution of Kin (Kin1-700-GFP) in control and homozygous *capu^EY12344^* mutant oocytes. This truncated motor protein lacks all its tail domains, and is thus not autoinhibited and unable to transport cargo (Williams *et al*., 2014). In control oocytes, Kin1-700 predominantly localizes to the posterior of the cell, in a gradient of diminishing abundance from anterior to posterior (Figure 2A and D, and (Williams *et al*., 2014)). In stark contrast, Kin1-700 fails to accumulate at the posterior in *capu* mutant oocytes (Figure 2B and F, Movie S10), and is found instead on filamentous structures closely resembling MTs (compare Figure 2B to Figure 1L and Movie S10 to Movie S5). This result indicates that, in the absence of *capu*, Kin becomes strongly recruited to MTs. The same Kin behavior can be induced acutely by *ex vivo* treatment with F-actin destabilising drugs like Cytochalasin D (data now shown), indicating that the observed behavior of Kin is due to the lack of the actin mesh and not to potential developmental defects that might arise from the lack of *capu* during earlier stages of oogenesis. In fixed control cells, Kin1-700 accumulates to posterior regions of the cell (Figure 2D), where MTs are less abundant (Figure 2C), while the motor strongly co-localizes with MTs in fixed *capu* mutant oocytes (Figure 2E, F, and J-L). This observation suggests that the actin mesh alters the ability of Kin to become recruited to MTs.

To further understand the relationship between the actin mesh and the interaction of Kin with MTs, we monitored the localization of the motor in different *capu* mutant alleles. For this, we utilized three mutant alleles that harbor distinct point mutations in the FH2 domain, and that affect the *in vitro* actin assembly rate of Capu to different degress (*capu^L219^, capu^L201^* and *capu^38^*, in increasing severity) (Yoo *et al*., 2015). In order to exclude effects from potential second site mutations, we generated trans-heterozygous animals between these alleles and the strong hypomorphic mutation *capu^EY12344^* (Figure 2 and TableS1). In order to estimate the amount of cytoplasmic F-actin, we measured relative signal intensities in phalloidin-stained control and mutant oocytes (Figure 2M-Q). In agreement with the *in vitro* data (Yoo *et al*., 2015), *capu^L219^* oocytes displayed a mild decrease in signal intensities, followed by *capu^L201^* and *capu^38^*, resulting in the strongest reduction in fluorescence intensities (Figure 2Q). We next tested the localization of Kin1-700 in these three allelic combinations of *capu*, and found that in *capu^L219^* oocytes, Kin is found at the posterior region of the cell (Figure 2R), but also at small MT bundles (arrow in Figure 2R). This recruitment of Kin to MTs was further enhanced in *capu^L201^* and *capu^38^*, to levels similar to what we observed in the homozygous *capu^EY12344^* cells (Figure 2S and T, compare to Figure 2B). Thus, the strength of Kin recruitment to MTs seems to follow the strength of the defects in the actin mesh. It is important to note that the speed of cytoplasmic flows is equally increased in *capu^L219^, capu^L201^* and *capu^38^* oocytes ((Yoo *et al*., 2015) and data not shown). Taken together, this indicates that the amount of Kin recruited to MTs mainly depends on the amount of cytoplasmic F-actin. Together with our previous finding that a higher number of active Kin results in faster flows (Ganguly *et al*., 2012), these findings point towards a model by which the actin mesh is regulating the magnitude of flows by limiting the efficient recruitment of Kin to MTs.

### Cytoplasmic streaming is necessary for MTs to display a correct orientation

Our results so far show that Kin-dependent cytoplasmic flows need to be maintained at low speeds in st9 oocytes for MTs not to mis-orient. Consequently, we next asked whether cytoplasmic streaming might actually be necessary to sustain a proper organization of the MT network. In other words, is the motion direction of MT plus ends altered in st9 oocytes that lack Kin-dependent cytoplasmic streaming?

To address this question, we first analyzed MTs in oocytes lacking *kinesin heavy chain* (*khc*). The complete loss of the Kin motor unit (mutant allele *khc^27^*, from hereon called *khc_null_*) results in a fully penetrant absence of cytoplasmic flows (Figure S8)(Palacios and St Johnston, 2002; Serbus *et al*., 2005). Compared to control cells, MTs in *khc^null^* mutant oocytes display very little overall motion and appear rather ‘stiff’ (Figure 3A,A’, compared to Figure 1J,J’ and Movie S6). However, oocytes without Kin do not only lack cytoplasmic streaming, but also lack cargo transport towards the ‘posterior tip’ and display an aberrant actin mesh (Figure 3D and Figure S9) (Brendza et al., 2000; Drechsler et al., 2017). To test whether the altered bulk behavior of MTs is due to the lack of streaming or other aberrations in Kin mutant oocytes, we monitored MTs in st9 oocytes carrying distinct mutations in the Kin motor domain, resulting in a slower motor (two mutant alleles known as *khc^23^* and *khc^17^*, hereafter summarized as *khc^slow^*) (Brendza et al., 1999). St9 *khc^slow^* oocytes display a normal cytoplasmic actin mesh (Figure S9) and are able to transport a considerable amount of cargo towards the posterior (Figure 3E and F) (Serbus *et al*., 2005; Loiseau *et al*., 2010). Importantly, and identical to *khc^null^* cells, they lack any cytoplasmic advection at st9, allowing us to study the impact of flows on MT behavior in the presence of transport (Figure S8)(Serbus *et al*., 2005). Similar to *khc^null^* cells, MTs in *khc^slow^* oocytes appear stiff and no motion could be detected in kymographs (Figure 3B-C’ and Movie S6). These observations indicate that the altered motion of MTs in oocytes without Kin is indeed due to a lack of cytoplasmic streaming. Thus, Kin-dependent cytoplasmic advection is necessary for MTs to display a wild-type bulk motion.

We next analyzed EB1 directionality in *khc^slow^* oocytes (Figure 3G-K). As in control cells, we observed dynamic EB1 comets throughout the entire cytoplasm (Movie S7). However, in stark contrast, the distribution of orientation angles displayed a more focused bias towards the posterior of the cell (Figure 3I vs. Figure 1Q). Consequently, *khc^slow^* mutant oocytes displayed an increased posterior plus-tip bias, with 75% of EB1 signals directed towards posterior, compared to 66% in control cells (Figure 3J vs. Figure 1R and Table 1). Furthermore, 54% of these posterior-directed EB1 signals (the group consisting of 75% of all comets) displayed a ‘posterior tip’ orientation, compared to 40% in controls (Figure 3K vs. Figure 1S and Table 1). These findings indicate that the orientation of the growing MT plus ends in st9 oocytes does not only depend on nucleation or anchoring of minus ends, but also on the presence of cytoplasmic flows. In other words, advection is necessary for MTs to display a correct orientation.

To further investigate the impact of cytoplasmic flows and Kin activity on MT orientation, we analyzed the regional organization of the MT network along the A-P axis in control, *khc^null^* and *khc^slow^* oocytes. We divided each oocyte into an anterior and a posterior region (Figure 4A), and analyzed EB1 directionality in each of these two regions (Table 1). As previously shown, the posterior EB1 bias increases along the A-P axis of st9 oocytes (Figure 4B-D) (Parton et al., 2011). In the anterior region, we found 62% of signals directed towards posterior (Figure 4C), while this bias was increased to 74% in the posterior region (Figure 4D). As already demonstrated for the global posterior EB1 bias (Figure 4B), *khc^slow^* oocytes also showed a dramatic change of MT orientation along the A-P axis, with a 74%, posterior bias in the anterior region (Figure 4E vs. Figure 4C) and an even further increased 81% posterior bias in the posterior region (Figure 4F vs. Figure 4D). Additionally, the ratio of signals directed towards the ‘posterior tip’ (Figure 1C) in both anterior and posterior regions of the *khc^slow^* cell was substantially increased, compared to controls (Figure 4E and F, and Table 1). This clearly demonstrates that oocytes with a slower Kin, and thus without cytoplasmic streaming, display a stronger polarization of the entire MT network towards the posterior pole, a key region in the establishment of the embryonic body axis. While the lack of Kin (*khc^null^*) seemed to cause only minor defects in the global organization of the MT cytoskeleton (Figure 3L-Pand Figure 4B), the regional analysis of EB1 directionality in *khc^null^* oocytes revealed major differences compared to control cells. In the anterior region of *khc^null^* cells, we detected an unexpected drop of the posterior bias (56% compared to 66% in controls), which indicates that the complete lack of Kin does indeed affect MT network organization (Figure 4G vs. Figure 4C). It needs to be mentioned here that oocytes lacking Kin also fail to localize the nucleus, which in turn is associated with MT minus ends (Williams et al., 2014). Therefore, this observed MT behavior in *khc^null^* cells might primarily reflect the mis-localization of a certain subset of MTOCs in the cell. However, MTs in the posterior region of *khc^null^* cells displayed a slightly more focused growth towards the key ‘posterior tip’ (Figure 1C), similar to *khc^slow^* cells (Figure 4H vs. Figure 4D and F, right rosette panels, and Table 1).

Taken together, our data allow us to draw certain conclusions about the relationship of Kin-activity, cytoplasmic streaming, and the organization of MTs. Firstly, in st9 oocytes cytoplasmic flows need to be in a defined regime to ensure proper MT orientation. Secondly, in the presence of Kin-mediated transport, but in the absence of cytoplasmic flows (*khc^slow^* condition), there are more plus ends growing towards the posterior, suggesting that cytoplasmic streaming is necessary for MTs to polarize in the precise pattern observed in control cells. And thirdly, together with cytoplasmic flows, other Kin-mediated processes, such as cargo transport or nucleus anchoring affect the organization of the MT network, supporting previously published findings (Nieuwburg et al., 2017; Zimyanin et al., 2008).

### Reconstitution of cytoplasmic flows in st9 *khc^slow^* oocytes rescues MT orientation

Our data suggest that Kin-mediated cytoplasmic streaming is necessary for the MT network to completely adopt its wild-type organization in the st9 oocyte. To further investigate the connection between cytoplasmic advection and MT orientation, we analyzed EB1 directionality in *capu,khc^slow^* double mutant oocytes. With this experiment we aimed to test whether the re-establishment of cytoplasmic streaming in a mutant background is sufficient to rescue MT orientation.

It has been suspected that faster cytoplasmic flows in *capu* mutants can be slowed down again by introducing a *khc^slow^* mutation (Dahlgaard et al., 2007). Therefore, we generated a double mutant, harboring the alleles *capu^EY12344^* and *khc^17^*. To verify the functionality of this double mutant, we first investigated posterior cargo localization in fixed cells. As expected, ~85% of *capu,khc^slow^/+* cells (which are essentially *capu* mutants, Table S1) failed to correctly localize Staufen to the posterior pole of the cell (Figure 5A, n=20). In comparison, ~72% of *capu,khc^slow^* double mutant cells localized Staufen into a posterior crescent (n=25). However, the majority of oocytes that displayed Staufen in a crescent, also showed Staufen accumulation in posterior dots (n=11/18). This constitutes a phenotype that is usually associated with the *khc^slow^* alleles (Figure 5B vs. Figure 3E and F). Together, these data confirmed that our double mutant is comparable to the previously reported alleles (Dahlgaard et al., 2007), and that slow Kin is sufficient to rescue the major cargo localization defects seen in *capu* mutants. However, since cargo transport was not rescued to wild-type levels, and was instead found to be similar to *khc^slow^* oocytes (Loiseau et al., 2010), it is obvious that the decreased cargo transport efficiency of slow Kin cannot be rescued by re-introducing cytoplasmic streaming by the lack of *capu*. It is furthermore unclear whether this is due to a reduced translocation speed of slow Kin, or to a defect in cargo anchoring in *capu* mutant cells (Tanaka *et al*., 2011).

**Figure 5.**
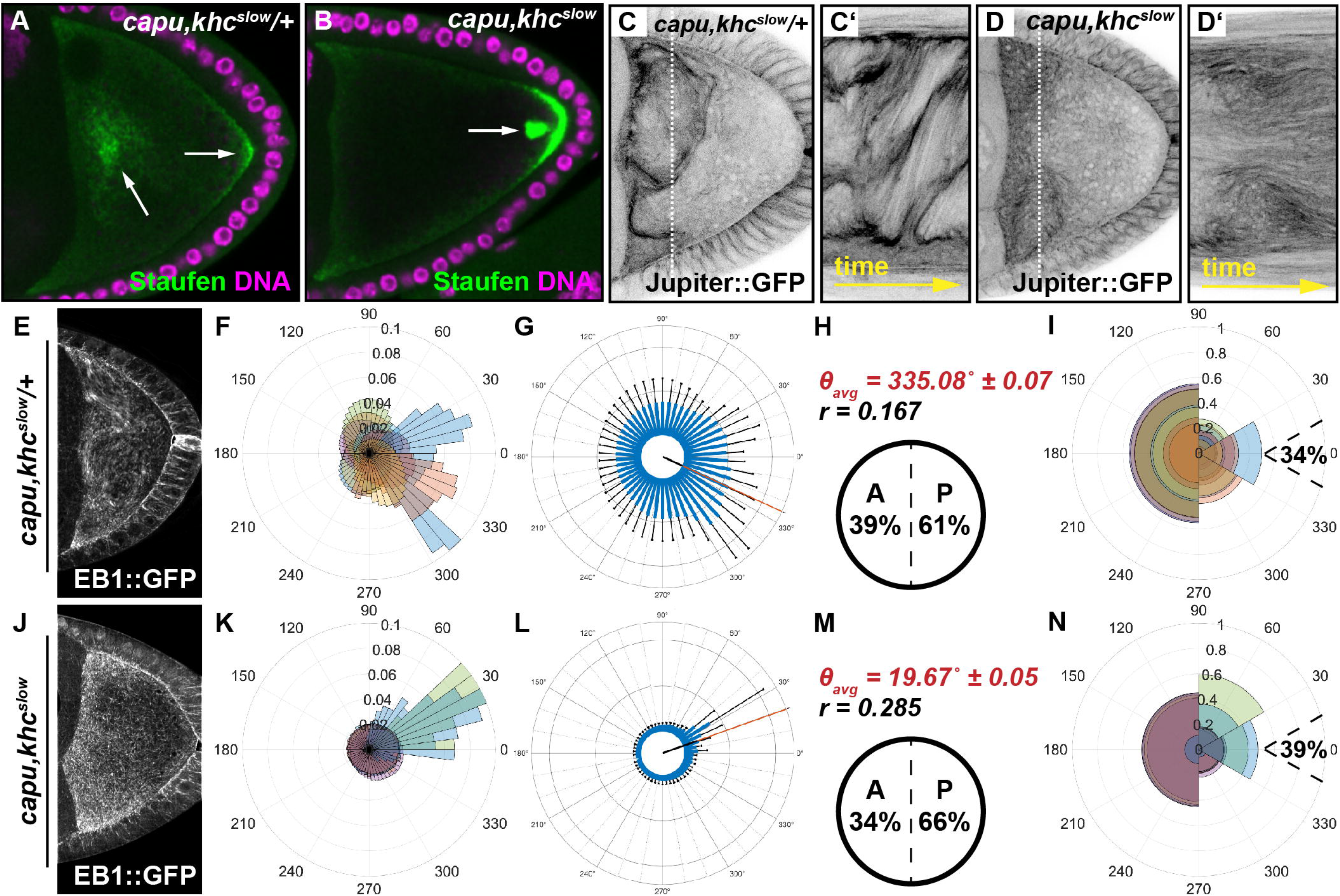
Cytoplasmic streaming constitutes a major contributor to MT orientation. **A,B)** Posterior cargo localization in *capu,khc^slow^/+* (A) and *capu,khc^slow^* double (B) mutant cells. While Staufen (green) partially localizes to the posterior of the cell in *capu,khc^slow^/+* oocytes (right arrow in A), the majority of the protein localizes in cytoplasmic clouds (left arrow in A). In *capu,khc^slow^* double mutant cells, Staufen localizes in a posterior crescent but also accumulates in dots close to the posterior (arrow in B). **C,D)** Still frames and kymographs of live *capu,khc^slow^/+* (C,C’) or *capu,khc^slow^* double (D,D’) mutant oocytes, expressing Jup. **E-N)** OF analysis of EB1 growth directionality in *capu,khc^slow^/+* (E-I, n=10) and *capu,khc^slow^* (J-N, n=8) oocytes.

Next we tested whether premature fast cytoplasmic flows in *capu* mutants are indeed slowed down again in *capu,khc^slow^* oocytes. As expected, and similar to *capu* single mutants, *capu,khc^slow^/+* oocytes displayed aberrant fast flows (Figure 5C, Movie S8). In contrast, cytoplasmic flows in *capu,khc^slow^* double mutant oocytes were indeed slowed down again and appeared similar to those observed in control cells (Figure S8). We then tested whether the rescue of cytoplasmic streaming in the double mutant also affected MT bulk movement. Compared to *capu,khc^slow^/+* oocytes, which displayed fast flows and consequently MT bundeling, the reduction of flow speeds in the double mutant caused a MT bulk movement similar to that observed in control cells (Figure 5C-D’ and Movie S8).

To finally test whether the re-establishment of cytoplasmic streaming in a *khc^slow^* background (or the reduction of fast flows in *capu* background) rescues the MT orientation defects observed in either single mutant, we analyzed EB1 directionality in *capu,khc^slow^/+* (Figure 5E-I) and *capu,khc^slow^* double mutant oocytes (Figure 5J-N and Movie S9). EB1 comets in *capu,khc^slow^/+* oocytes exhibited values that are similar to what we observed in *capu* mutants (Figure 1T-X), suggesting that the heterozygous presence of a *khc^slow^* mutant chromosome does not substantially affect the *capu* mutant phenotype (Movie S9, 61% of EB1 signals pointing towards posterior, and 34% of this 61% are growing towards the ‘posterior tip’ in *capu,khc^slow^/+* cells, Figure 5F-I and Table 1). Conversely, EB1 comets in *capu,khc^slow^* double mutant cells displayed a posterior motion bias indistinguishable from that of control cells (Movie S9, 66% posterior bias, Figure 5M vs. Figure 1R and Table 1). Finally, also the fraction of all posterior comets pointing towards the ‘posterior tip’ was found to be similar in *capu,khc^slow^* double mutant cells (39%) and controls (40%) (Figure 5N vs. Figure 1S and Table 1), further strengthening the idea that cytoplasmic streaming constitutes a substantial contributor to a correct MT plus end focussing, and thus MT organization, in the oocyte.

In summary, our results now clearly demonstrate that in the absence of *capu* and the actin mesh, a slower Kin motor protein is sufficient to restore the correct regime of cytoplasmic flows, resulting in the correct polarization of the MT cytoskeleton, and an almost restored transport of cargo towards the posterior. Therefore, it seems that the actin mesh is essential in oocytes that exhibit normal Kin-mediated transport, likely by regulating the amount of Kin recruited to MTs, in order to ensure the proper regulation of cytoplasmic streaming, which in turn is an important contributor to the observed MT orientation.

## DISCUSSION

Due to its large size (80-100 μm along the A-P axis) and the fact that MT minus ends are nucleated and anchored along the antero-lateral cortices of the cell, the *Drosophila* st9 oocyte is arguably one of the most challenging *in vivo* systems in which to analyze the dynamic behavior of MTs. The use of confocal microscopy, together with OF based analysis tool described in this study, allowed us to quantify the motion direction of plus ends within a 2D focal plane along the entire length of this cell. Together with others, the present study supports the existence of a complex mechanical and/or biochemical relationship between motion of cytoplasmic components (whether by advection or by transport), cytoplasmic F-actin and MTs in the *Drosophila* oocyte. This holds true for our model system, but is likely transferable to many other cell types. For example, experimental data and modeling of cytoplasmic motion in Characean algae (where cytoplasmic flows are acto-myosin dependent) suggests that paralle actin cables and ordered streaming patterns self-organize in an interdependent manner (Foissner and Wasteneys, 2000; Woodhouse and Goldstein, 2013).

Kin-mediated cargo transport through a highly viscous medium, such as the cytoplasm, inevitably induces bulk motion of such medium, which will cause lateral displacement forces on MT filaments and thus induce a feedback on their spatial orientation. As a consequence, cytoplasmic streaming needs to be kept in a defined speed regime and at a biased random pattern in order for the MT network to properly polarize. In the absence of the actin mesh (as in *capu* or *spire* mutants), there is an increased persistence and speed of flows, resulting in parallel alignment and bundling of MTs, as well as strong defects on MT orientation. The actin mesh - which itself requires Kin activity for its proper organization (Drechsler et al., 2017) - is required for the maintenance of this correct regime of advection.

With our work, we show that our image analysis approach - based on a rigorous image analysis framework - is able to infer and quantify directionality of EB1 motion from confocal image sequences. The use of an efficient iterative optimisation algorithm enabled us to analyze entire image sequences at once, as compared to a frame-by-frame analysis. In comparison to more commonly used techniques, such as particle tracking or PIV, OF allows us to perform motion estimation on pixel level in challenging circumstances, such as low signal-to-noise ratios and small particle size. Most importantly, this image analysis approach can handle standard confocal microscopy data and the method does not require demanding imaging techniques or costly computing hardware. Moreover, the image analysis depends only on a few parameters that can be easily adjusted. However, two limitations of the developed methodology need to be pointed out. Firstly, due to the use of a variational framework for the image analysis, both the denoised sequences and the estimated velocities exhibit a loss of contrast, leading to underestimated speeds of EB1 comets. This is particularly due to the temporal regularization required in both steps to overcome the above-mentioned challenges. Secondly, our statistical analyses of directions of EB1 comets are based on velocities computed for all pixels within each segmented oocyte. We are aware that not every image pixel portrays exactly one EB1 comet. Furthermore, EB1 comets in the oocyte are subjected to different forces. In summary however, our results demonstrate that our approach is able to monitor general plus end direction in most, if not all, cell types and thus constitutes an efficient and reliable analytical framework for MT polarity studies. Therefore, OF is absolutely suitable to efficiently describe the orientation of MTs in crowded intracellular environments.

It is unknown how the actin mesh slows down Kin-dependent cytoplasmic flows or how it might affect MT organization. It has been proposed that the presence of a viscoelastic actin network can increase the effective viscosity of the cytoplasm, and counteract the viscous drag of cargo transport by Kin (Quinlan, 2016). Furthermore, actin-MT crosslinking proteins have been demonstrated to allow a potent crosstalk between both filament species (Preciado Lopez *et al*., 2014), presumably coordinating cytoskeletal organization *in vivo*. However, our data from *capu,khc^slow^* double mutant oocytes, which do not form an actin mesh, suggest that the mesh itself is not essential to allow a correct MT orientation when Kin is slower. In this ‘artificial’ mutant situation, the correct regime of flows seems sufficient to allow proper MT network organization. It is thus possible that the actin mesh regulates the activity of Kin more directly, for example by tethering the motor to its filaments (directly or indirectly through cargo). Such model is indeed supported by our finding that Kin becomes efficiently recruited to MTs when the actin mesh is absent (Movie S10 and Figure 2). This observation could explain how the mesh regulates not only flows, but also the higher degree of MT bundling, as a larger number of Kin molecules recruited to MTs is likely to result in faster flows (Ganguly *et al*., 2012). Furthermore, this could also result in a higher degree of effective crosslinking of adjacent filaments (Lu et al., 2016). However, the recruitment of Kin to MTs depends on various regulatory mechanisms, including interactions with cargo and MT-associated proteins like Ensconsin (Metivier *et al*., 2019). For this reason we used the truncated form Khc1-700 in our analyses. Whether, and how, the actin mesh influences the interaction of full length Kin with MTs - in the presence of other MT-binding regulatory mechanisms - needs to be further addressed in the future. Nevertheless, our findings indicate a novel potential mechanism of how the actin and MT cytoskeletons can regulate each other.

Our data establish cytoplasmic streaming in st9 oocytes as a contributing factor for the correct organization of the MT cytoskeleton. If - and how - a stronger posterior polarization of the MT network may affect cargo delivery to the posterior is unknown. Tracking *oskar* mRNA particles in *khc^slow^* oocytes did not reveal a stronger orientation bias of cargo movement (Zimyanin *et al*., 2008). However, it is unclear whether trajectories of *oskar* transport always reflect the organization of the underlying MT network, and whether all MTs would be equally used by slow Kin motor/cargo complexes.

Previous studies suggested that the pattern of nucleation and minus end anchoring along the cortex are sufficient to organize the MT network and to allow correct cargo transport to define the A-P axis of the animal. Consequently, mutant oocytes harboring defects in nucleation and/or anchoring of minus ends also display polarity defects (Doerflinger et al., 2006; Nashchekin et al., 2016). Furthermore, mathematical modeling suggested that cytoplasmic streaming at mid-oogenesis is negligible to explain the correct localization of posterior cargoes like *oskar* mRNA (Khuc Trong *et al*., 2015). However, our study demonstrates that cytoplasmic streaming is involved in the orientation of MT plus ends, and we conclude that the localization of the minus ends alone is not sufficient to define the precise overall organization of the network. This was most obvious in *khc^slow^* oocytes, which in our hands lack cytoplasmic streaming entirely, and displayed an increased posterior orientation MT bias (Figure 3). The advantage of analyzing slow Kin mutants is that other Kin-dependent processes, like cargo transport and formation of the actin mesh, do take place. Despite showing only mild defects in the distribution of developmental determinants, *khc^slow^* mutant oocytes fail to give rise to a healthy offspring, suggesting that oocyte polarity is affected (Moua *et al*., 2011). Consistently, we found an increased posterior bias of EB1 signals in *khc^slow^* mutant oocytes, strongly suggesting that the lack of cytoplasmic streaming was causative for this observation.

In summary, the combination of various forces produced within living cells demands a complex set of biochemical and biomechanical regulatory mechanisms for cytoskeletal networks to organize correctly. Our results show that, in *Drosophila* st9 oocytes, advection by cytoplasmic streaming contributes to the polarization of MTs by affecting the directional motion of MT plus ends. All these observations further stress the need for a combination of different experimental approaches in order to fully understand the dynamic organization of cytoskeletons, from simplified *in vitro* systems to complex *in vivo* situations.

## METHODS

### Fly stocks and genetics

Flies were kept at standard corn meal agar and raised at room temperature (21°C). Detailed genotypes of all fly stocks can be found in Table S1. Germ line clones for the analysis of *khc* mutant alleles have been induced by the FLP/FRT ovoD system (Chou and Perrimon, 1996). Germline clones in Figure 5 were identified by the absence of nuclear GFP in germline cells.

### Live imaging

Female flies of the desired genotypes were collected and fattened on dry yeast for 12-16 h prior to imaging. Ovaries were dissected in a small drop of halocarbon oil (Voltalef S10, VWR) on a glass coverslip and single egg chambers were separated using fine tungsten needles. Images were acquired on a Leica SP5 inverted confocal microscope, using a 40×/1.3 Oil DIC Plan-Neofluar (Jup) or a 100×/1.4 Oil DIC objective (EB1). Signals were detected using a Leica HyD Hybrid Detector. For MT bulk movement, a single plane from the middle of the oocyte was imaged at a scan speed of 100 Hz and at an image resolution of 1,024 × 1,024 pixels (corresponding to one image every 10.4 s). For EB1 imaging the oocyte was fitted and oriented within a 512 x 256 pixels frame and a single plane image was taken every 0.65 s. Image sequences of at least 100 frames (65 s) were taken, inspected visually, and bleach corrected using Fiji (Schindelin *et al*., 2012). In all experiments the middle of the oocyte was defined as the focal plane in which the oocyte had its largest area.

### Immunostainings

Egg chambers were dissected in PBS+0.1% Tween20 and fixed in 10% formaldehyde in PBS+0.1% Tween20 for 10 min. Fixed ovaries were incubated with anti-αTubulin (MAB1864, Sigma-Aldrich, cloneYL1/2, 1:100) or anti-Staufen (gift from. D. St Johnston, 1:3,000) in PBS+2% Tween20 over night at 4°C. After four consecutive washes Alexa-coupled secondary antibodies (1:100) were incubated for two hours at room temperature. Native fluorescence of GFP was imaged without amplification. Images were acquired on a Leica SP5 inverted confocal microscope, using a 40×/1.3 Oil DIC Plan-Neofluar objective. For phalloidin staining, egg chambers where dissected and fixed in 10% methanol free formaldehyde in PBS, containing 0.1% Tween-20 (PBT0.1), for a maximum of 10 min. Fixed cells were washed 4x in PBT0.1 and incubated with 1□μM TRITC-coupled phalloidin (in PBT0.1, Sigma-Aldrich) over night at 4□°C. The stained samples were washed 4x in PBT0.1, mounted, and imaged imediatly under identical conditions, using a Zeiss LSM800 equipped with a 40×/1.3 Oil DIC Plan-Neofluar objective. Normalized fluorescence intensities were acquired using FiJi.

### Image denoising and optical flow based motion estimation

Motion analysis of the recorded two-dimensional image sequences was performed using a two-step procedure. The first step aimed to remove noise contamination from the unprocessed sequences, while the goal of the second step was to estimate motion in terms of displacement vector fields from the so-improved sequences. As noise may significantly disturb the motion estimation (Ruhnau et al., 2005; Burger *et al*., 2018), this two-step procedure is crucial for obtaining robust estimation results. In the first step, we recovered from each recorded image sequence *u^δ^* an improved version *u* by solving a variational image denoising problem with spatio-temporal regularization. It reads

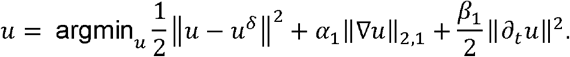

While the first two terms on the right-hand side resemble standard image denoising with total variation regularization (Rudin L.I, 1992) in space, the third term connects subsequent frames by penalising temporal changes within the recovered solution. As EB1 comets typically appear in several subsequent frames at similar positions, it allows to effectively remove randomly distributed noise from a sequence (see Figure 2D,E, Movie S2, and SI). Here, *α,β* >0 are regularization parameters that balance the three terms and need to be chosen appropriately. Moreover, the norms (as defined in equation S21) are taken over the entire image sequence.

The result *u* served as input to the motion estimation step, in which we estimated a displacement vector field 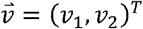 by solving

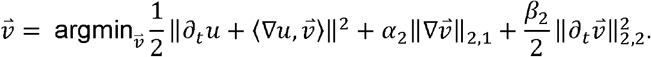

Here, 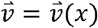 is defined for every pixel *x* of the image *u* and is an estimate of the motion that *u* experiences at *x* from one image frame to the subsequent one in time. That is, the vector 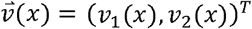 points in the direction of motion at *x*, and 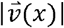 is an estimate of the speed of the motion at *x*. The first term on the right-hand side in the above equation aims to approximately solve the optical flow equation (Horn and Schunck, 1981), while the second and third terms incorporate spatio-temporal regularization of the vector-valued unknown (for further details see SI). The use of the vector-valued total variation allows for spatial discontinuities in the displacement vector field. We found that in both steps the temporal regularization was key and the analysis of individual frames did not yield satisfactory results. Both minimisation problems were approximately solved using the primal-dual hybrid gradient method (Chambolle, 2011) and graphics-processing unit (GPU) acceleration. An in-depth description of both models, their numerical solution, and parameter choices as well as implementation details can be found in the SI.

## ACKNOWLEDGEMENTS

We are greatful to Dr. Fabio Giavazzi (Milan) for help with the PIV analysis in some of our experiments. We thank Drs D. St Johnston and I. Davis for reagents, and M. Wayland for assistance with imaging. We also thank Drs Lena Frerking and Sujoy Ganguly for fruitful discussions during the initial phase of the project, and Drs MD Martin Bermudo and D. St Johnston for comments on the manuscript and discussions.

MD and IMP were supported by the BBSRC, the University of Cambridge and Queen Mary University of London. MD was also supported by an Isaac Newton Trust fellowship and acknowledges funding by the DFG/SFB944 (‘Physiology and Dynamics of Cellular Microcompartments’), the University of Osanbrück and the State of Lower Saxony. LFL and CBS acknowledge support from the Leverhulme Trust (‘Breaking the non-convexity barrier’), EPSRC (grant No. EP/M00483X/1), the EPSRC Centre (No. EP/N014588/1), the RISE projects CHiPS and NoMADS, the Cantab Capital Institute for the Mathematics of Information, and the Alan Turing Institute. LAK was supported by Queen Mary University of London. MB and HD acknowledge support by the European Research Council (EU FP7-ERC Consolidator Grant No. 615216 LifeInverse). We gratefully acknowledge the support of NVIDIA Corporation with the donation of the Quadro P6000 GPU used for this research.

## AUTHOR CONTRIBUTIONS

MD designed and performed research, analyzed data, and wrote the paper. IMP designed research, analyzed data, and wrote the paper. LAK performed research and analyzed data. LFL and CBS developed new analytic tools, analyzed the data, and wrote the paper. HD and MB contributed analytic tools.

## COMPETING INTERESTS

The authors declare no competing financial interests.

## DATA AVAILABILITY

The source code of our implementation and of the data analysis is available online (https://doi.org/10.5281/zenodo.2573254). All relevant data and the computational results are available from the corresponding authors upon request.

## Notes

https://doi.org/10.5281/zenodo.2573254

## REFERENCES

Akhmanova, A., and Steinmetz, M.O. (2015). Control of microtubule organization and dynamics: two ends in the limelight. Nat Rev Mol Cell Biol 16, 711–726.

Amat, F., Myers, E.W., and Keller, P.J. (2013). Fast and robust optical flow for time-lapse microscopy using super-voxels. Bioinformatics 29, 373–380.

Baker, S., Scharstein D, Lewis J.P, Roth S, Black M.J, and Szeliski R. (2011). A database and evaluation methodology for optical flow. Int. J. Comput. Vision 92(1), 1–31.

Boquet-Pujadas, A., Lecomte, T., Manich, M., Thibeaux, R., Labruyere, E., Guillen, N., Olivo-Marin, J.C., and Dufour, A.C. (2017). BioFlow: a non-invasive, image-based method to measure speed, pressure and forces inside living cells. Sci Rep 7, 9178.

Boric, K., Orio, P., Vieville, T., and Whitlock, K. (2013). Quantitative analysis of cell migration using optical flow. PLoS One 8, e69574.

Brendza, R.P., Serbus, L.R., Duffy, J.B., and Saxton, W.M. (2000). A function for Kinesin I in the posterior transport of *oskar* mRNA and Staufen protein. Science 289, 2120–2122.

Burger, M., Dirks, H., and Schönlieb, C.-B. (2018). A Variational Model for Joint Motion Estimation and Image Reconstruction. Siam Journal on Imaging Sciences 11, 94–128.

Chambolle, A.P., T.. (2011). A first-order primal-dual algorithm for convex problems with applications to imaging.. J. Math. Imaging Vis. 40, 120–145.

Chou, T.B., and Perrimon, N. (1996). The autosomal FLP-DFS technique for generating germline mosaics in *Drosophila melanogaster*. Genetics 144, 1673–1679.

Dahlgaard, K., Raposo, A.A., Niccoli, T., and St Johnston, D. (2007). Capu and Spire assemble a cytoplasmic actin mesh that maintains microtubule organization in the *Drosophila* Oocyte. Dev Cell 13, 539–553.

Delpiano, J., Jara J, Scheer J, Ramírez O.A, Ruiz-del-Solar J, and S., H. (2012). Performance of optical flow techniques for motion analysis of fluorescent point signals in confocal microscopy. Machine Vision and Applications 23, 675–689.

Doerflinger, H., Benton, R., Torres, I.L., Zwart, M.F., and St Johnston, D. (2006). *Drosophila* anterior-posterior polarity requires actin-dependent PAR-1 recruitment to the oocyte posterior. Curr Biol 16, 1090–1095.

Drechsler, M., Giavazzi, F., Cerbino, R., and Palacios, I.M. (2017). Active diffusion and advection in *Drosophila* oocytes result from the interplay of actin and microtubules. Nat Commun 8, 1520.

Fassier, C., Freal, A., Gasmi, L., Delphin, C., Ten Martin, D., De Gois, S., Tambalo, M., Bosc, C., Mailly, P., Revenu, C., Peris, L., Bolte, S., Schneider-Maunoury, S., Houart, C., Nothias, F., Larcher, J.C., Andrieux, A., and Hazan, J. (2018). Motor axon navigation relies on Fidgetin-like 1-driven microtubule plus end dynamics. J Cell Biol 217, 1719–1738.

Foissner, I., and Wasteneys, G.O. (2000). Microtubule disassembly enhances reversible cytochalasin-dependent disruption of actin bundles in characean internodes. Protoplasm 214, 33–44.

Forrest, K.M., and Gavis, E.R. (2003). Live imaging of endogenous RNA reveals a diffusion and entrapment mechanism for nanos mRNA localization in *Drosophila*. Curr Biol 13, 1159–1168.

Frerking, L., Burger, M., Vestweber, D., and C., B. (2014). TGV-based flow estimation for 4D leukocyte transmigration. A. K. Louis, S. Arridge, and B. Rundell, editors, Proceedings of the Inverse Problems from Theory to Applications Conference, 79–83.

Ganguly, S., Williams, L.S., Palacios, I.M., and Goldstein, R.E. (2012). Cytoplasmic streaming in *Drosophila* oocytes varies with kinesin activity and correlates with the microtubule cytoskeleton architecture. Proc Natl Acad Sci U S A 109, 15109–15114.

Gonzalez-Reyes, A., Elliott, H., and St Johnston, D. (1995). Polarization of both major body axes in *Drosophila* by gurken-torpedo signalling. Nature 375, 654–658.

Guo, v.d.V., and Zhou. (2014). Red blood cell tracking using optical flow methods. IEEE J. Biomed. Health Inform 18(3), 991–998.

Huang, Y., Hao, L., Li, H., Liu, Z., and Wang, P. (2017). Quantitative Analysis of Intracellular Motility Based on Optical Flow Model. J Healthc Eng 2017.

Hurd, T.R., Herrmann, B., Sauerwald, J., Sanny, J., Grosch, M., and Lehmann, R. (2016). Long Oskar Controls Mitochondrial Inheritance in *Drosophila melanogaster*. Dev Cell 39, 560–571.

Januschke, J., Gervais, L., Dass, S., Kaltschmidt, J.A., Lopez-Schier, H., Johnston, D.S., Brand, A.H., Roth, S., and Guichet, A. (2002). Polar transport in the *Drosophila* oocyte requires Dynein and Kinesin I cooperation. Curr Biol 12, 1971–1981.

Jiang, K., Toedt, G., Montenegro Gouveia, S., Davey, N.E., Hua, S., van der Vaart, B., Grigoriev, I., Larsen, J., Pedersen, L.B., Bezstarosti, K., Lince-Faria, M., Demmers, J., Steinmetz, M.O., Gibson, T.J., and Akhmanova, A. (2012). A Proteome-wide screen for mammalian SxIP motif-containing microtubule plus-end tracking proteins. Curr Biol 22, 1800–1807.

Kent, I.A., Rane, P.S., Dickinson, R.B., Ladd, A.J., and Lele, T.P. (2016). Transient Pinning and Pulling: A Mechanism for Bending Microtubules. PLoS One 11, e0151322.

Khuc Trong, P., Doerflinger, H., Dunkel, J., St Johnston, D., and Goldstein, R.E. (2015). Cortical microtubule nucleation can organise the cytoskeleton of *Drosophila* oocytes to define the anteroposterior axis. Elife 4.

Kodama, A., Karakesisoglou, I., Wong, E., Vaezi, A., and Fuchs, E. (2003). ACF7. An essential integrator of microtubule dynamics. Cell 115, 343–354.

Loiseau, P., Davies, T., Williams, L.S., Mishima, M., and Palacios, I.M. (2010). *Drosophila* PAT1 is required for Kinesin-1 to transport cargo and to maximize its motility. Development 137, 2763–2772.

Lu, W., Lakonishok, M., Serpinskaya, A.S., Kirchenbuechler, D., Ling, S.C., and Gelfand, V.I. (2018). Ooplasmic flow cooperates with transport and anchorage in *Drosophila* oocyte posterior determination. J Cell Biol 217, 3497–3511.

Lu, W., Winding, M., Lakonishok, M., Wildonger, J., and Gelfand, V.I. (2016). Microtubule-microtubule sliding by kinesin-1 is essential for normal cytoplasmic streaming in *Drosophila* oocytes. Proc Natl Acad Sci U S A 113, E4995–5004.

Metivier, M., Monroy, B.Y., Gallaud, E., Caous, R., Pascal, A., Richard-Parpaillon, L., Guichet, A., Ori-McKenney, K.M., and Giet, R. (2019). Dual control of Kinesin-1 recruitment to microtubules by Ensconsin in *Drosophila* neuroblasts and oocytes. Development 146.

Mitchison, T., and Kirschner, M. (1984). Dynamic instability of microtubule growth. Nature 312, 237–242.

Monteith, C.E., Brunner, M.E., Djagaeva, I., Bielecki, A.M., Deutsch, J.M., and Saxton, W.M. (2016). A Mechanism for Cytoplasmic Streaming: Kinesin-Driven Alignment of Microtubules and Fast Fluid Flows. Biophys J 110, 2053–2065.

Moua, P., Fullerton, D., Serbus, L.R., Warrior, R., and Saxton, W.M. (2011). Kinesin-1 tail autoregulation and microtubule-binding regions function in saltatory transport but not ooplasmic streaming. Development.

Nashchekin, D., Fernandes, A.R., and St Johnston, D. (2016). Patronin/Shot Cortical Foci Assemble the Noncentrosomal Microtubule Array that Specifies the *Drosophila* Anterior-Posterior Axis. Dev Cell 38, 61–72.

Nieuwburg, R., Nashchekin, D., Jakobs, M., Carter, A.P., Khuc Trong, P., Goldstein, R.E., and St Johnston, D. (2017). Localised dynactin protects growing microtubules to deliver oskar mRNA to the posterior cortex of the *Drosophila* oocyte. Elife 6.

Palacios, I.M., and St Johnston, D. (2002). Kinesin light chain-independent function of the Kinesin heavy chain in cytoplasmic streaming and posterior localisation in the *Drosophila* oocyte. Development 129, 5473–5485.

Parton, R.M., Hamilton, R.S., Ball, G., Yang, L., Cullen, C.F., Lu, W., Ohkura, H., and Davis, I. (2011). A PAR-1-dependent orientation gradient of dynamic microtubules directs posterior cargo transport in the *Drosophila* oocyte. J Cell Biol 194, 121–135.

Preciado Lopez, M., Huber, F., Grigoriev, I., Steinmetz, M.O., Akhmanova, A., Koenderink, G.H., and Dogterom, M. (2014). Actin-microtubule coordination at growing microtubule ends. Nat Commun 5, 4778.

Purro, S.A., Ciani, L., Hoyos-Flight, M., Stamatakou, E., Siomou, E., and Salinas, P.C. (2008). Wnt regulates axon behavior through changes in microtubule growth directionality: a new role for adenomatous polyposis coli. J Neurosci 28, 8644–8654.

Quinlan, M.E. (2013). Direct interaction between two actin nucleators is required in *Drosophila* oogenesis. Development 140, 4417–4425.

Quinlan, M.E. (2016). Cytoplasmic Streaming in the Drosophila Oocyte. Annu Rev Cell Dev Biol 32, 173–195.

Roth, S., Neuman-Silberberg, F.S., Barcelo, G., and Schupbach, T. (1995). *cornichon* and the EGF receptor signaling process are necessary for both anterior-posterior and dorsal-ventral pattern formation in *Drosophila*. Cell 81, 967–978.

Rudin L.I, O.S., and Fatemi E. (1992). Nonlinear total variation based noise removal algorithms. Physica D: Nonlinear Phenomena 60, 259–268.

Ruhnau et al. (2005). Variational optical flow estimation for particle image velocimetry. Expe. Fluids 38(1), 21–32.

Schindelin, J., Arganda-Carreras, I., Frise, E., Kaynig, V., Longair, M., Pietzsch, T., Preibisch, S., Rueden, C., Saalfeld, S., Schmid, B., Tinevez, J.Y., White, D.J., Hartenstein, V., Eliceiri, K., Tomancak, P., and Cardona, A. (2012). Fiji: an open-source platform for biological-image analysis. Nat Methods 9, 676–682.

Serbus, L.R., Cha, B.J., Theurkauf, W.E., and Saxton, W.M. (2005). Dynein and the actin cytoskeleton control kinesin-driven cytoplasmic streaming in *Drosophila* oocytes. Development 132, 3743–3752.

Tanaka, T., Kato, Y., Matsuda, K., Hanyu-Nakamura, K., and Nakamura, A. (2011). *Drosophila* Mon2 couples Oskar-induced endocytosis with actin remodeling for cortical anchorage of the germ plasm. Development 138, 2523–2532.

Theurkauf, W.E. (1994). Premature microtubule-dependent cytoplasmic streaming in *cappuccino* and *spire* mutant oocytes. Science 265, 2093–2096.

Vig, D.K., Hamby, A.E., and Wolgemuth, C.W. (2016). On the Quantification of Cellular Velocity Fields. Biophys J 110, 1469–1475.

Williams, L.S., Ganguly, S., Loiseau, P., Ng, B.F., and Palacios, I.M. (2014). The auto-inhibitory domain and ATP-independent microtubule-binding region of Kinesin heavy chain are major functional domains for transport in the *Drosophila* germline. Development 141, 176–186.

Woodhouse, F.G., and Goldstein, R.E. (2013). Cytoplasmic streaming in plant cells emerges naturally by microfilament self-organization. Proc Natl Acad Sci U S A 110, 14132–14137.

Yoo, H., Roth-Johnson, E.A., Bor, B., and Quinlan, M.E. (2015). *Drosophila* Cappuccino alleles provide insight into formin mechanism and role in oogenesis. Mol Biol Cell 26, 1875–1886.

Zimyanin, V.L., Belaya, K., Pecreaux, J., Gilchrist, M.J., Clark, A., Davis, I., and St Johnston, D. (2008). *In vivo* imaging of *oskar* mRN A transport reveals the mechanism of posterior localization. Cell 134, 843–853.

