## Supplementary Information for "Optical flow analysis reveals that Kinesin-mediated advection impacts on the orientation of microtubules in the *Drosophila* oocyte"

### Supplementary Methods

In this section, we discuss the image analysis and computational methods used in this study in more detail. First, we state the variational image denoising and motion estimation models used in our two-step approach. Second, we explain the efficient numerical solution of these models and give all required implementation details. Third, we discuss the appropriate choices of regularisation parameters used in the models. Finally, we briefly outline the workflow of our analysis of growth directionality of EB1-labelled comets and explain the computation of the used statistical quantities.

#### Motion Analysis based on Variational Optical Flow

Optical flow (OF) refers to the displacement vector field that describes the apparent motion in a sequence of images [14]. Variational OF methods constitute a popular framework for reliable and efficient dense motion estimation with a wide range of applications. See e.g. [4] for a recent survey and e.g. [1] for an introduction.

One of the main challenges when solving for the displacement vector field in the confocal image sequences at hand is the combination of a very low signal-to-noise ratio and the relatively small size of the fluorescently labelled EB1 comets. As a remedy, we apply a preceding image denoising step that effectively eliminates noise, but is able to preserve fluorescence response of EB1-labelled comets up to a known loss of contrast [19].

To this end, we consider a discrete (scalar) image sequence that is arranged as a vector $u^\delta \in \mathbb{R}^M$ with $M = mnt$, where $n$ and $m$ denote the number of pixels in vertical, respectively, horizontal direction, and $t$ denotes the number of frames. The finite-dimensional image denoising problem then reads

$$\min_u \frac{1}{2}\|u - u^\delta\|^2 + \alpha_1 \|D_x u\|_{2,1} + \frac{\beta_1}{2}\|D_t u\|^2, \tag{S1}$$

where $u \in \mathbb{R}^M$ is the unknown solution, and $D_x : \mathbb{R}^M \to \mathbb{R}^{M\times 2}$ and $D_t : \mathbb{R}^M \to \mathbb{R}^M$ are finite difference approximations of the spatial and the temporal gradient operators, respectively.

The first two terms in (S1) constitute a standard model for variational image denoising of videos, see e.g. [1]. The first term is a data discrepancy term, which ensures that the solution $u$ is close to the original sequence $u^\delta$, and the second term is a discrete (isotropic) approximation of the (scalar) *total* *variation* regularisation [15]. We use first-order forward differences with step size one and zero-Neumann boundary conditions, see e.g. [9], and define

$$\|D_x u\|_{2,1} = \sum_{j=1}^{M} \sqrt{(D_x u)_{j,1}^2 + (D_x u)_{j,2}^2}.$$

In addition, the third term in (S1) penalises changes over time. The regularisation parameters $\alpha_1, \beta_1 > 0$ control the tradeoff between the three terms and need to be chosen carefully and in dependence of the noise level [16]. We assume that the noise level is similar in all recorded image sequences. In our experiments we found that these parameters significantly impact the subsequent motion estimation. The functional in (S1) enforces temporal regularity of the reconstructed image sequence and thereby effectively removes noise, as it is—in contrast to the fluorescence response of the EB1-labelled comets—assumed to be randomly distributed in space and time.

In the second step, we estimate displacement vector fields from the improved sequence $u$. For $\mathbf{v} = (v_1, v_2) \in \mathbb{R}^{N\times 2}$ with $N = mn(t-1)$, the finite-dimensional minimisation problem reads

$$\min_{\mathbf{v}} \ \frac{1}{2}\|\partial_t u + \langle \nabla u, \mathbf{v}\rangle\|^2 + \alpha_2 \|\mathbf{D}_x \mathbf{v}\|_{2,1} + \frac{\beta_2}{2}\|\mathbf{D}_t \mathbf{v}\|_{2,2}^2, \tag{S2}$$

and incorporates spatio-temporal regularisation of the unknown. Here, the first term enforces the *optical flow equation* [14] to be solved approximately. Moreover, $\partial_t u \in \mathbb{R}^N$ and $\nabla u \in \mathbb{R}^{N\times 2}$ refer to discrete approximations of temporal and spatial partial derivatives, respectively, and the scalar product in the first term is taken along rows.

The matrices $\mathbf{D}_x : \mathbb{R}^{N\times 2} \to \mathbb{R}^{N\times 4}$ and $\mathbf{D}_t : \mathbb{R}^{N\times 2} \to \mathbb{R}^{N\times 2}$ again are finite difference approximations of the spatial and temporal gradient of a vector-valued quantity, respectively. They are given by

$$\mathbf{D}_x \mathbf{v} = (D_x v_1, D_x v_2) \qquad \text{and} \qquad \mathbf{D}_t \mathbf{v} = (D_t v_1, D_t v_2).$$

As an (isotropic) discrete approximation of the *vector total variation*, see e.g. [6], we use

$$\|\mathbf{D}_x \mathbf{v}\|_{2,1} = \sum_{j=1}^{N} \sqrt{(D_x v_1)_{j,1}^2 + (D_x v_1)_{j,2}^2 + (D_x v_2)_{j,1}^2 + (D_x v_2)_{j,2}^2},$$

see e.g. [9, Sec. 7.5]. In the terminology of [11], this corresponds to the *collaborative total variation* using the $\ell^{2,2,1}(\mathrm{der}, \mathrm{col}, \mathrm{pix})$ norm, see [11, Sec. 1.2 and Table 1], and $\|\cdot\|_{2,2}$ is the matrix $\ell^{2,2}$ norm. We highlight that this definition of the total variation for vector-valued functions couples its components. For other choices see e.g. the references in [11]. Again, $\alpha_2, \beta_2 > 0$ are regularisation parameters that need to be chosen appropriately.

In our experiments we found that the temporal regularisation in (S1) and (S2) was crucial, and a frame by frame analysis of the image sequences proved insufficient due to the abovementioned challenges.

#### Numerical Solution

In order to solve problems (S1) and (S2) numerically, we solve the corresponding saddle-point problems up to sufficient accuracy. The above minimisation problems are of the general form

$$\min_{x\in X} \ f(Kx) + g(x). \tag{S3}$$

Here, $K : X \to Y$ is a continuous linear operator between finite-dimensional real Hilbert spaces $X$ and $Y$, and both $f : Y \to [0, +\infty]$ and $g : X \to [0, +\infty]$ are proper, convex, and lower-semicontinuous functions. The corresponding saddle point formulation reads

$$\max_{y\in Y} \inf_{x\in X} \ \langle y, Kx\rangle - f^*(y) + g(x), \tag{S4}$$

where $f^* : Y^* \to [0, +\infty]$ is the Legendre–Fenchel conjugate of $f$ and $Y^*$ denotes the dual space of $Y$. Problem (S4) can be solved by means of the primal-dual hybrid gradient (PDHG) algorithm [8], which consists of iterating

$$\begin{cases} x^{(k+1)} = \mathrm{prox}_{\tau g}\Big(x^{(k)} - \tau K^* y^{(k)}\Big), \\ y^{(k+1)} = \mathrm{prox}_{\sigma f^*}\Big(y^{(k)} + \sigma K\big(2x^{(k+1)} - x^{(k)}\big)\Big), \end{cases} \tag{S5}$$

up to the desired accuracy for given initial data $x^{(0)}, y^{(0)}$. Here, $\tau, \sigma > 0$ are parameters and $\mathrm{prox}(\cdot)$ denotes the proximal map, given by

$$\mathrm{prox}_J(\tilde{x}) = \arg\min_x \frac{1}{2}\|x - \tilde{x}\|^2 + J(x),$$

with $J$ being a proper, convex, and lower-semicontinuous function. See e.g. [9, Sec. 3.4].

Assuming that (S4) admits solutions, so-called saddle points, and $\tau\sigma\|K\|^2 < 1$ is satisfied, then iteration (S5) converges to a saddle point at the rate $O(1/k)$ [8, Thm. 1]. Here, $\|K\|$ denotes the operator norm of $K$. We refer e.g. to [9] for further details.

##### Image Denoising

The problem (S1) can be written in the form of (S3) with the operator $K : \mathbb{R}^M \to \mathbb{R}^{M\times 3}$ given by $Ku = (D_x u, D_t u)$ and

$$g(u) = \frac{1}{2}\|u - u^\delta\|^2, \qquad f(\mathbf{p}, q) = \alpha_1\|\mathbf{p}\|_{2,1} + \frac{\beta_1}{2}\|q\|^2. \tag{S6}$$

Here, $\mathbf{p} \in \mathbb{R}^{M\times 2}$ and $q \in \mathbb{R}^M$.[1] Since the function $f$ is separable, its Legendre–Fenchel conjugate is

$$f^*(\mathbf{p}, q) = \delta_{\{\|\cdot\|_{2,\infty}\le\alpha_1\}}(\mathbf{p}) + \frac{1}{2\beta_1}\|q\|^2, \tag{S7}$$

see e.g. [3, Thm. 4.12]. Here, $\delta_{\{\|\cdot\|_{2,\infty}\le\alpha_1\}}$ is the indicator function of the dual ball of size $\alpha_1$. It is defined as

$$\delta_{\{\|\cdot\|_{2,\infty}\le\alpha_1\}}(\mathbf{p}) = \begin{cases} 0 & \text{if } \|\mathbf{p}_j\| \le \alpha_1 \text{ for all } j, \\ +\infty & \text{else,} \end{cases}$$

where the index $j$ refers to the $j$-th row of a matrix. A corresponding saddle point formulation is then

$$\min_u \max_{\mathbf{p},q} \langle D_x u, \mathbf{p}\rangle + \langle D_t u, q\rangle + g(u) - \delta_{\{\|\cdot\|_{2,\infty}\le\alpha_1\}}(\mathbf{p}) - \frac{1}{2\beta_1}\|q\|^2. \tag{S8}$$

The proximal mappings in (S5) with respect to $g$ and $f^*$ can be computed in a straightforward manner. For the function $g$ in (S6) it is given by

$$\mathrm{prox}_{\frac{\tau}{2}\|\cdot - u^\delta\|^2}(\tilde{u}) = \frac{\tilde{u} + \tau u^\delta}{1 + \tau}.$$

Since $f^*$ is separable, the proximal mapping of (S7) can be computed separately for each dual variable $\mathbf{p}$ and $q$, see e.g. [3, Thm. 6.6]. They are given by the (pixelwise) orthogonal projection onto the $\|\cdot\|_{2,\infty}$-ball of size $\alpha_1$ [8, Eq. 4.23], that is

$$\hat{\mathbf{p}} = \Pi_{\{\|\cdot\|_{2,\infty}\le\alpha_1\}}(\tilde{\mathbf{p}}) \Leftrightarrow \hat{\mathbf{p}}_j = \frac{\tilde{\mathbf{p}}_j}{\max\{1, \alpha_1^{-1}\|\tilde{\mathbf{p}}_j\|\}},$$

and by

$$\mathrm{prox}_{\frac{\sigma}{2\beta_1}\|\cdot\|^2}(\tilde{q}) = \frac{\beta_1\tilde{q}}{\sigma + \beta_1}.$$

##### Motion Estimation

Similar to before, the problem (S2) can be written in the form of (S3) with the operator $K : \mathbb{R}^{N\times 2} \to \mathbb{R}^{N\times 6}$ given by $K\mathbf{v} = (\mathbf{D}_x\mathbf{v}, \mathbf{D}_t\mathbf{v})$ and

$$g(\mathbf{v}) = \frac{1}{2}\|\partial_t u + \langle\nabla u, \mathbf{v}\rangle\|^2, \qquad f(\mathbf{p}, \mathbf{q}) = \alpha_2\|\mathbf{p}\|_{2,1} + \frac{\beta_2}{2}\|\mathbf{q}\|_{2,2}^2. \tag{S9}$$

Here, $\mathbf{p} \in \mathbb{R}^{N\times 4}$ and $\mathbf{q} \in \mathbb{R}^{N\times 2}$. As before, $f$ is a separable function in $\mathbf{p}$ and $\mathbf{q}$, and we have

$$f^*(\mathbf{p}, \mathbf{q}) = \delta_{\{\|\cdot\|_{2,\infty}\le\alpha_2\}}(\mathbf{p}) + \frac{1}{2\beta_2}\|\mathbf{q}\|_{2,2}^2. \tag{S10}$$

[1] Even though uncommon in the literature, we will refrain from introducing new variables and instead use the same for both $f$ and $f^*$ for simplicity, since the dimensions are the same.

A corresponding saddle point formulation is

$$\min_{\mathbf{v}} \max_{\mathbf{p},\mathbf{q}} \langle \mathbf{D}_x \mathbf{v}, \mathbf{p} \rangle + \langle \mathbf{D}_t \mathbf{v}, \mathbf{q} \rangle + g(\mathbf{v}) - \delta_{\{\|\cdot\|_{2,\infty} \leq \alpha_2\}}(\mathbf{p}) - \frac{1}{2\beta_2} \|\mathbf{q}\|_{2,2}^2. \tag{S11}$$

It remains to state the proximal mappings required for iterating (S5). For the function $g$ in (S9) it can be computed as the solution of a linear system of two equations for each (pixel) $j \in \{1, \ldots, N\}$, see e.g. [10, Chap. 4.5.2]. It is given by

$$\hat{\mathbf{v}} = \operatorname{prox}_{\frac{\tau}{2}\|\partial_t u + \langle \nabla u, \cdot \rangle\|^2}(\tilde{\mathbf{v}}) \Leftrightarrow \hat{\mathbf{v}}_j \text{ solves } \left(\mathrm{Id} + \tau (\nabla u)_j^\top (\nabla u)_j\right) \hat{\mathbf{v}}_j^\top = \tilde{\mathbf{v}}_j^\top - \tau (\nabla u)_j^\top (\partial_t u)_j,$$

where $\mathrm{Id}$ is the identity matrix of size two. Moreover, since $f^*$ again is separable with respect to $\mathbf{p}$ and $\mathbf{q}$, the proximal map with respect to (S10) can be computed separately for each variable. Similar to before, they are given by

$$\hat{\mathbf{p}} = \Pi_{\{\|\cdot\|_{2,\infty} \leq \alpha_2\}}(\tilde{\mathbf{p}}) \Leftrightarrow \hat{\mathbf{p}}_j = \frac{\tilde{\mathbf{p}}_j}{\max\{1, \alpha_2^{-1} \|\tilde{\mathbf{p}}_j\|\}}$$

and by

$$\operatorname{prox}_{\frac{\sigma}{2\beta_2}\|\cdot\|_{2,2}^2}(\tilde{\mathbf{q}}) = \frac{\beta_2 \tilde{\mathbf{q}}}{\sigma + \beta_2}.$$

##### Implementation Details

We implemented algorithm (S5) in MATLAB using built-in graphics-processing unit (GPU) acceleration. Computations were performed on an Intel Xeon E5-2630 v4 $2.2\,\mathrm{GHz}$ server equipped with $128\,\mathrm{GB}$ RAM and an NVIDIA Quadro P6000 GPU featuring $24\,\mathrm{GB}$ of memory.

A typical two-dimensional image sequence $u^\delta$ has $m = 256$ horizontal, respectively $n = 512$ vertical pixels, and $t = 100$ frames (although some sequences are longer and have $t = 200$ or even $t = 400$ frames). Before processing, the intensity values of each sequence were scaled to the interval $[0, 1]$. The total number of unknowns in problems (S1) and (S2) for a typical sequence are $M \approx 13 \cdot 10^6$ and $N \approx 26 \cdot 10^6$, respectively, plus the number of corresponding dual variables. A typical sequence with 100 frames thus results in approximately 13 million computed (displacement) vectors.

As a termination criterion in algorithm (S5) we used the primal-dual residual, see e.g. [13]. By defining

$$\begin{aligned} p^{(k+1)} &:= \frac{1}{\tau}\big(x^{(k)} - x^{(k+1)}\big) - K^*\big(y^{(k)} - y^{(k+1)}\big), \\ d^{(k+1)} &:= \frac{1}{\sigma}\big(y^{(k)} - y^{(k+1)}\big) - K\big(x^{(k)} - x^{(k+1)}\big), \end{aligned}$$

we say that convergence in terms of the primal-dual residual is achieved if

$$\frac{1}{N} \sum_j \left| p_j^{(k+1)} \right| + \frac{1}{N} \sum_j \left| d_j^{(k+1)} \right| < \varepsilon. \tag{S12}$$

Here, the index $j$ runs over all entries of the corresponding vector or matrix. In our experiments we set $\varepsilon := 10^{-6}$ and terminated algorithm (S5) as soon as (S12) was satisfied. For computational efficiency, (S12) was checked only every 100 iterations for problem (S8) and every 500 iterations for problem (S11).

A straightforward calculation along the lines of [7, Thm. 3.1] gives $\|K\|^2 \leq 12$ for both formulations. This can easily be confirmed numerically by applying the power method, see e.g. [18], to $K^*K$, since $\|K\|^2 = \|K^*\|^2 = \|K^*K\|$. However, we found experimentally that, for both problems, algorithm (S5) converged in terms of the primal-dual residual with parameters set to $\tau, \sigma := 1/\sqrt{8}$.

The total number of processed image sequences amounted to 57. On average, the algorithm was terminated after $788.14 \pm 85.53$ ($95\,\%$ confidence interval) iterations for the image denoising and after $7788.14 \pm 492.44$ iterations for the motion estimation, which resulted in an average computation time per sequence of $1.18 \pm 0.16$ minutes and $27.37 \pm 4.22$ minutes, respectively, using the GPU. The overall processing time per sequence was $28.55 \pm 4.29$ minutes on average. In total, the computational analysis took approximately 28 hours.

The source code of our implementation and of the data analysis is available online.[2]

[2] https://doi.org/10.5281/zenodo.2573254

#### Estimation of Regularisation Parameters

In order to select appropriate regularisation parameters in problems (S1) and (S2), we chose one representative sequence and computed results for the first ten frames for varying parameters $\alpha_j, \beta_j$, where $j \in \{1, 2\}$. Supplementary Fig. S1 shows the chosen test sequence. Observe the high noise contamination and the small size of the fluorescently labelled EB1 comets.

We solved problems (S1) and (S2) based on this test sequence for all parameter combinations

$$(\alpha_1, \beta_1, \alpha_2, \beta_2) \in \{0.005, 0.01, 0.05\} \times \{0.05, 0.1, 0.75\} \times \{0.0005, 0.001, 0.05\} \times \{0.001, 0.005, 0.01\}.$$

Supplementary Fig. S1 shows the solution $u$ of the denoising step for one particular parameter setting. To be precise, it shows the variable $u$ of an approximate solution of the saddle point problem (S8). In Suppl. Fig. S2, we illustrate results for various parameter configurations. We only show the first frame of each result. The effect of the parameter $\alpha_1$ is clearly visible and choosing it too large results in large piecewise constant and undesirable patches (see Suppl. Fig. S2, bottom row). The results for varying parameter $\beta_1$ are omitted, as its effect is difficult to visualise.

In Suppl. Fig. S3, we show results for the motion estimation step for fixed parameters $\alpha_2, \beta_2$ but varying parameters $\alpha_1, \beta_1$ for the denoising problem. The same parameters as in Suppl. Fig. S2 were used to demonstrate the impact of the denoising step on the result of the motion estimation. Again, we display the approximation $\mathbf{v}$ of a saddle point of problem (S11). In Suppl. Fig. S3, we use a standard colour-coding to display the computed velocities [2]. For a vector $\mathbf{v}_j^\top \in \mathbb{R}^2$ at a pixel $j$, the displayed colour is determined by the orientation of the velocity, while the shown intensity depends on its magnitude. The colour-coding is indicated at the boundary of the results. In order to increase the contrast, the velocities were scaled before visualisation by adjusting the histogram of their magnitudes slightly.

It is evident that varying the regularisation parameters of the denoising step significantly impacts the result of the motion estimation. Clearly, a certain level of regularisation is required. Choosing the parameter $\alpha_1$ too large results in a loss of information and the subsequent motion estimation is not able to identify individual EB1-labelled comet movement, see red arrows in Suppl. Fig. S3. Conversely, choosing parameter $\beta_1$ too small, results in the appearance of false signals in the velocity fields, see black arrows in Suppl. Fig. S3.

In order to determine parameter settings that capture EB1 motion accurately, we determined flow fields for the first 10 frames of a single image sequence by testing each individual of the 81 possible combinations of regularisation parameters $\alpha_j, \beta_j$, where $j \in \{1, 2\}$. Based on the resulting vector fields, the vast majority of these combinations could be excluded by visual analysis, since they resulted in a complete loss of motion signals, or in noisy vector fields indicating motion in areas of the cell that were clearly void of EB1 comets. The final parameter setting that we used was then chosen based on visual comparison of the computed vector field with the appearance of EB1 signals in the image series. Since a ground truth (e.g. EB1 tracks or clean image sequences) is not available for our data sets, testing a finite number of combinations for the regularisation parameters was necessary to estimate suitable values for the further analysis.

The final parameter setting used to analyse the sequences is highlighted in Suppl. Fig. S2 with a red and in Suppl. Fig. S3 with a black frame, respectively. Let us mention that, since we obtained our image sequences from a single imaging plane, the estimated velocity is the apparent 2D motion of fluorescent particles and does not account for movement orthogonal to the imaging plane.

#### Computational and Statistical Analysis

In Suppl. Fig. S4 we show results of the two-step image analysis for one selected control cell. The denoised sequence shows a visually improved signal-to-noise ratio (see also Suppl. Movie 2) and the colour-coded displacement fields indicate movement of bright spots very well. Moreover, in Suppl. Fig. S5 we show the mean velocities (over time) within each masked oocyte for all analysed datasets. The result shown in Suppl. Fig. S4 corresponds to the top left result in Suppl. Fig. S5.

As a verification of the approach and the selected regularisation parameters, we examined the arithmetic mean velocity (over time) $\bar{\mathbf{v}}$ in a control region around the follicular cells in the posterior of the cells, see Suppl. Fig. S6(A)–(C). In this region, the growth of microtubules is known to take place predominantly outwards in radial direction (i.e. to the right). The growth direction of the majority of EB1 comets is very well captured, see Suppl. Fig. S6(C) and Suppl. Movie 3.

In addition, in Suppl. Fig. S6(D) we visualise the direction of velocities $\mathbf{v}$ in this control region with the help of a rose diagram (angular histogram). It is created by computing the map $\mathbf{v} \mapsto (\theta, \rho)$ that converts velocities in Cartesian representation to their representation in polar coordinates with $\theta$ being the angle (direction) and $\rho$ the radius (speed). The angular histogram is then created by plotting a circular histogram of the directions $\theta$ using 50 bins. The height of each bar is determined by the relative number of angles $\theta$ in each bin. Note that, in this representation the magnitude of $\mathbf{v}$ is not taken into account and that the shown polar histogram also includes velocities outside the follicular epithelium (faint regions in the top and bottom right corners in Suppl. Fig. S6(C)), which we neglect for simplicity.

Both the mean velocity, shown in Suppl. Fig. S6(C), and the angular histogram of $\theta$, shown in Suppl. Fig. S6(D), provide strong evidence that the apparent (average) growth of microtubules in outward radial direction is captured well by the two-step procedure with the selected parameters. In Suppl. Fig. S7 we illustrate the mean velocities in the control regions for all the analysed image sequences. The results are widely consistent and show outward growth of EB1 comets. Moreover, cell boundaries of follicle cells are respected and in some cases trajectories of individual EB1 comets are visible, cf. Suppl. Movie 3.

The computational workflow for the analysis of the captured sequence was as follows. For each image sequence:

1. Solve image denoising problem (S1) to obtain $u$.
2. Solve motion estimation problem (S2) based on $u$ to obtain displacements $\mathbf{v}$.
3. Scale displacements according to pixel size $\Delta x$ and the time interval $\Delta t = 0.65\,\mathrm{s}$ between consecutive frames to obtain approximate velocities $\mathbf{v} := \mathbf{v}\Delta x/\Delta t$ (in $\mu\mathrm{m}/s$).
4. Create a hand-drawn segmentation mask of the oocyte using Fiji [17]; see also Fig. 2(H).
5. Restrict velocities $\mathbf{v}$ to the mask.
6. Compute mean flow $\bar{\mathbf{v}}$ for visualisation purpose, angles $\theta$ and speeds $\rho$, and output statistics.

The scaling step in 3. is due to the chosen discretisation of the spatial and temporal gradient operators. It is however insubstantial for the analysis of the directionality.

As the main focus of our analysis was the directionality of the movements of EB1-labelled comets, we computed descriptive statistics of the angles of the estimated velocities. These are circular quantities and require appropriate methods, for instance, when computing a mean angle. We refer to e.g. [5, 12] for more details on the following concepts.

Given a vector of angles $\theta \in \mathbb{R}^N$ of estimated velocities (e.g. of one cell), we consider the mean resultant vector $\mathbf{v}_{\mathrm{avg}} \in \mathbb{R}^2$, which is given by

$$\mathbf{v}_{\mathrm{avg}} = \frac{1}{N} \sum_{j=1}^{N} \begin{pmatrix} \cos\theta_j \\ \sin\theta_j \end{pmatrix}. \tag{S13}$$

Switching to the complex plane, it can easily be computed as $\mathbf{v}_{\mathrm{avg}} = \frac{1}{N}\sum_j \mathrm{e}^{\mathrm{i}\theta_j}$, where $\mathrm{i}$ is the imaginary unit. Then, the mean angular direction $\theta_{\mathrm{avg}}$ and the resultant vector length $r$ are given by $\theta_{\mathrm{avg}} = \mathrm{angle}(\mathbf{v}_{\mathrm{avg}})$ and $r = \|\mathbf{v}_{\mathrm{avg}}\|$, respectively. The circular variance is related to $r$ and is defined as $S = 1 - r$.

In Suppl. Fig. S6(F)–(I), we illustrate the analysis of a selected cell. It shows the angular histogram for directions $\theta$ using 50 bins for multiple cells, the mean angular direction, and the mean resultant vector for the velocities within the segmented oocytes.

### References


[1] G. Aubert and P. Kornprobst. *Mathematical problems in image processing*, volume 147 of *Applied Mathematical Sciences*. Springer, New York, 2 edition, 2006.

[2] S. Baker, D. Scharstein, J. P. Lewis, S. Roth, M. J. Black, and R. Szeliski. A database and evaluation methodology for optical flow. *Int. J. Comput. Vision*, 92(1):1–31, 2011.

[3] A. Beck. *First-Order Methods in Optimization*. Society for Industrial and Applied Mathematics, 2017.

[4] F. Becker, S. Petra, and Ch. Schnörr. Optical flow. In O. Scherzer, editor, *Handbook of Mathematical Methods in Imaging*, pages 1956–2004. Springer New York, 2015.

[5] P. Berens. CircStat: A MATLAB toolbox for circular statistics. *J. Stat. Softw.*, 31(10), 2009.

[6] X. Bresson and T. Chan. Fast dual minimization of the vectorial total variation norm and applications to color image processing. *Inverse Probl. Imaging*, 2(4):455–484, 2008.

[7] A. Chambolle. An algorithm for total variation minimization and applications. *J. Math. Imaging Vision*, 20(1–2):89–97, 2004.

[8] A. Chambolle and T. Pock. A first-order primal-dual algorithm for convex problems with applications to imaging. *J. Math. Imaging Vision*, 40(1):120–145, 2011.

[9] A. Chambolle and T. Pock. An introduction to continuous optimization for imaging. *Acta Numer.*, 25:161–319, 2016.

[10] H. Dirks. *Variational Methods for Joint Motion Estimation and Image Reconstruction*. PhD thesis, Institute for Computational and Applied Mathematics, University of Münster, Germany, 2015.

[11] J. Duran, M. Moeller, C. Sbert, and D. Cremers. Collaborative total variation: A general framework for vectorial TV models. *SIAM J. Imaging Sciences*, 9(1):116–151, 2016.

[12] N. I. Fisher. *Statistical Analysis of Circular Data*. Cambridge University Press, 1993.

[13] T. Goldstein, M. Li, X. Yuan, E. Esser, and R. Baraniuk. Adaptive primal-dual hybrid gradient methods for saddle-point problems. Technical report, 2015.

[14] B. K. P. Horn and B. G. Schunck. Determining optical flow. *Artif. Intell.*, 17:185–203, 1981.

[15] L. I. Rudin, S. Osher, and E. Fatemi. Nonlinear total variation based noise removal algorithms. *Phys. D*, 60(1–4):259–268, 1992.

[16] O. Scherzer, M. Grasmair, H. Grossauer, M. Haltmeier, and F. Lenzen. *Variational Methods in Imaging*. Number 167 in Applied Mathematical Sciences. Springer, New York, 2009.

[17] J. Schindelin, I. Arganda-Carreras, E. Frise, V. Kaynig, M. Longair, T. Pietzsch, S. Preibisch, C. Rueden, S. Saalfeld, B. Schmid, J.-Y. Tinevez, D. J. White, V. Hartenstein, K. Eliceiri, P. Tomancak, and A. Cardona. Fiji: an open-source platform for biological-image analysis. *Nature*, 9(7):676–682, 2012.

[18] G. Strang. *Linear algebra and its applications*. Brooks/Cole/Cengage, fourth edition, 2006.

[19] D. Strong and T. Chan. Edge-preserving and scale-dependent properties of total variation regularization. *Inverse Probl.*, 19(6):165–187, 2003.

### Supplementary Figures

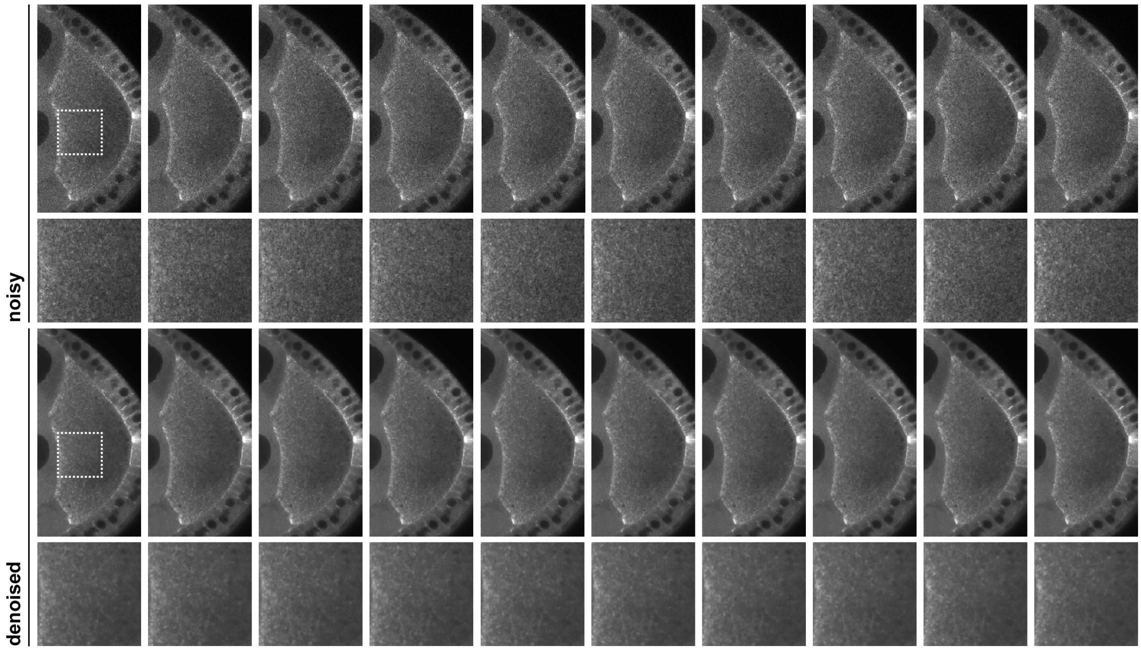


**Figure S1 - Image denoising using a specific choice of regularisation parameters.** The top rows show the ten frames of the unprocessed and noisy test sequence $u^\delta$ used for estimating the regularisation parameters (magnified views in the second row are taken from the area indicated by the dashed box). The bottom panels show the denoised sequence $u$ (magnified views in the bottom row are taken from the area indicated by the dashed box), obtained by approximately solving saddle point formulation (S8) using the parameters $\alpha_1 = 0.005$ and $\beta_1 = 0.75$.

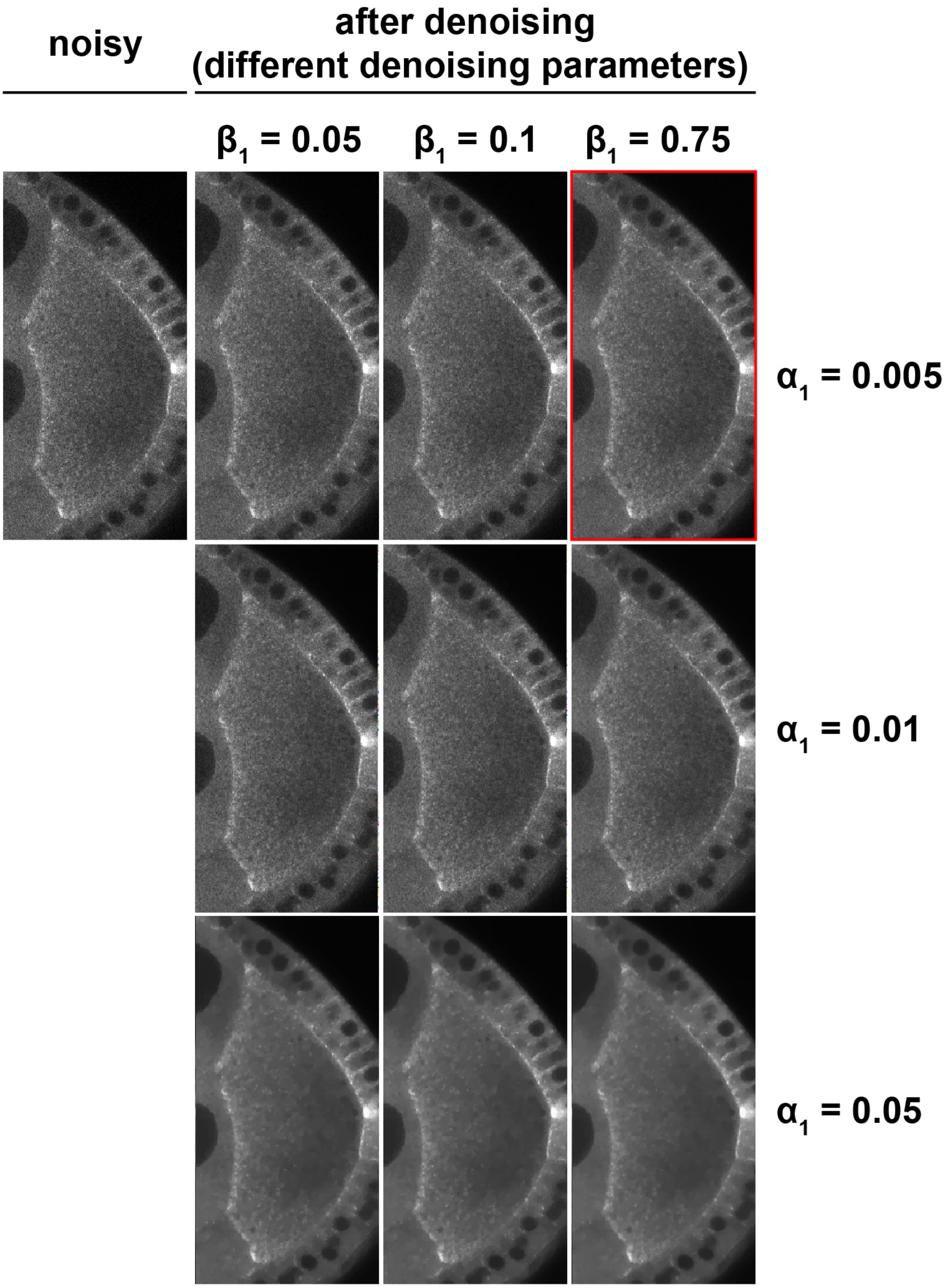


**Figure S2 - Image denoising step using different regularisation parameters.** Depicted are the first frame of the unprocessed test sequence $u^\delta$ (upper left, as shown in Suppl. Fig. S1) and the first frame of the denoised test sequence $u$ for varying parameters $\alpha_1$ and $\beta_1$ (as indicated). The red frame (upper right image) indicates the parameter setting that was used to analyse the sequences (same as in Suppl. Fig. S1).

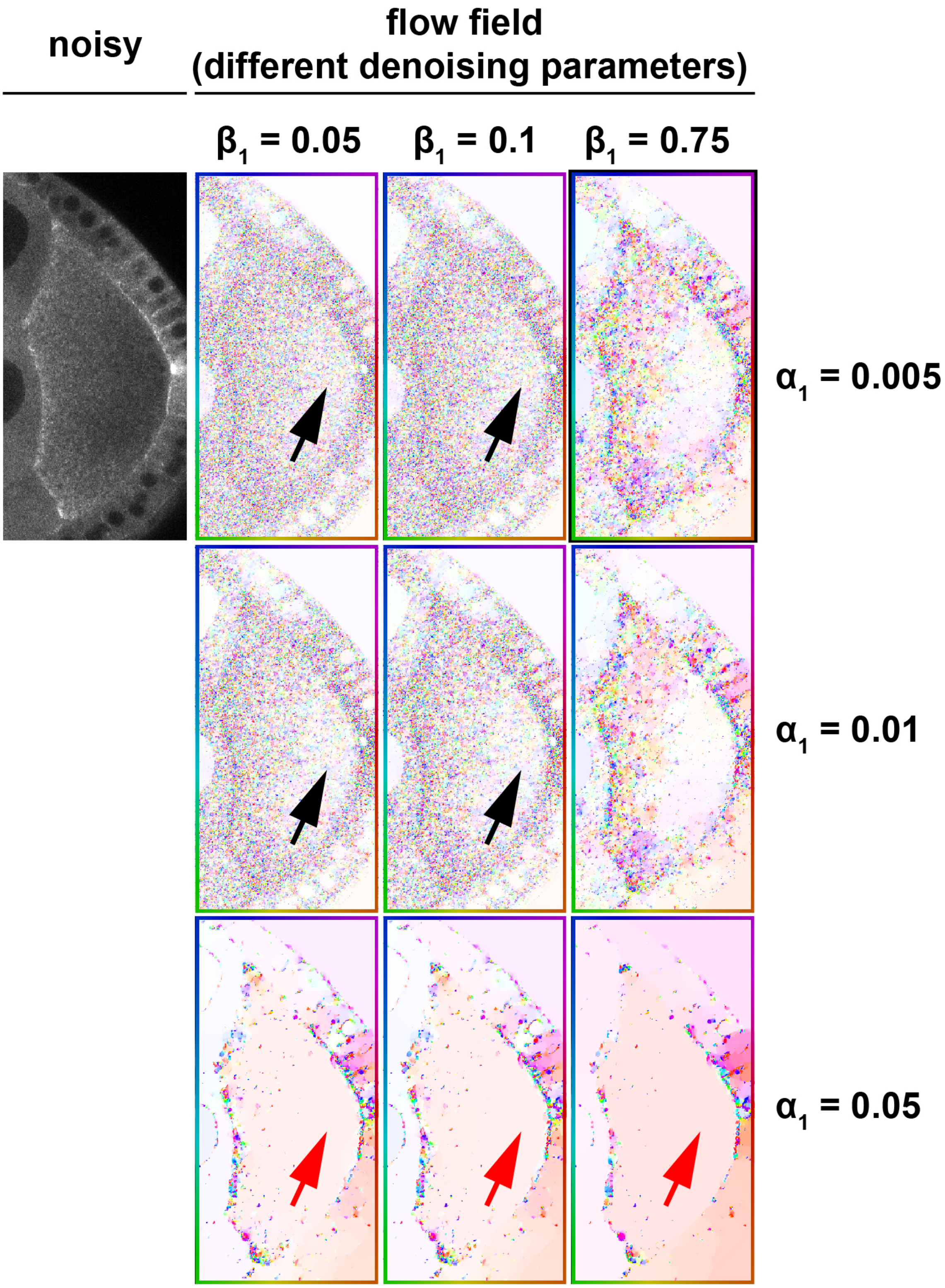


**Figure S3 - Effect of denoising parameter choice on OF motion estimation.** Velocities $\mathbf{v}$ estimated by approximately solving (S11) for the denoised sequences for different regularisation parameter choices (as indicated). Shown is the colour-coded displacement field $\mathbf{v}$ between the first pair of frames. For all results, the same parameter setting for the motion estimation was used ($\alpha_2 = 0.0005$ and $\beta_2 = 0.005$), while the parameters for the denoising step were varied as indicated. Black arrows mark flow fields in which the parameter settings resulted in unsatisfying, i.e. too noisy, signals after the motion estimation. The red arrows label flow fields that lost signal due to too much regularisation. The black frame (upper right image) indicates the final parameter setting that was used to analyse the sequences.

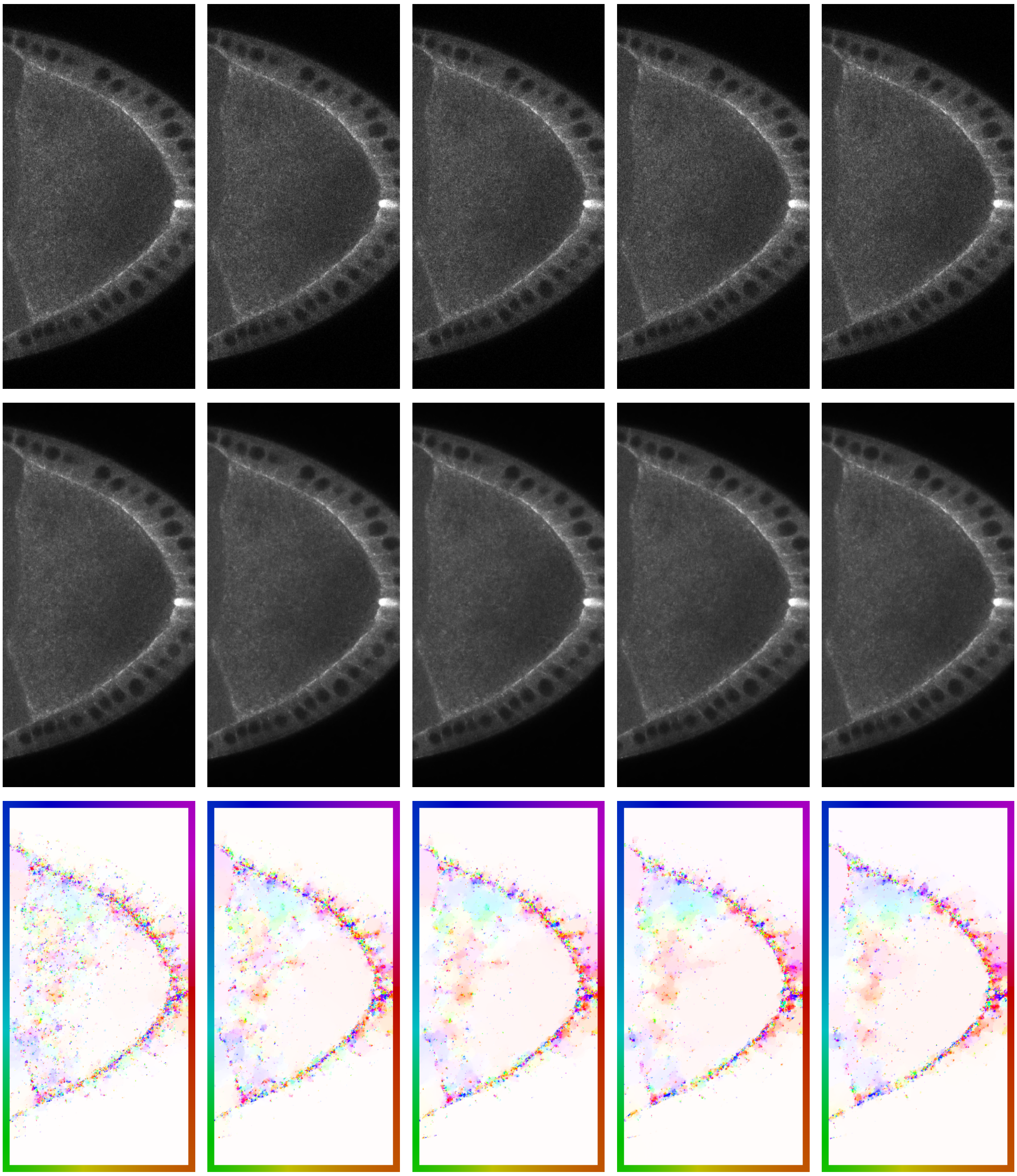

**Figure S4 - Computational workflow for one image sequence.** The figure shows the first five frames of the original noisy image sequence $u^\delta$ (top row), the denoised sequence $u$ (middle row), and the first five colour-coded velocity fields $\mathbf{v}$ (bottom row) for a selected cell.

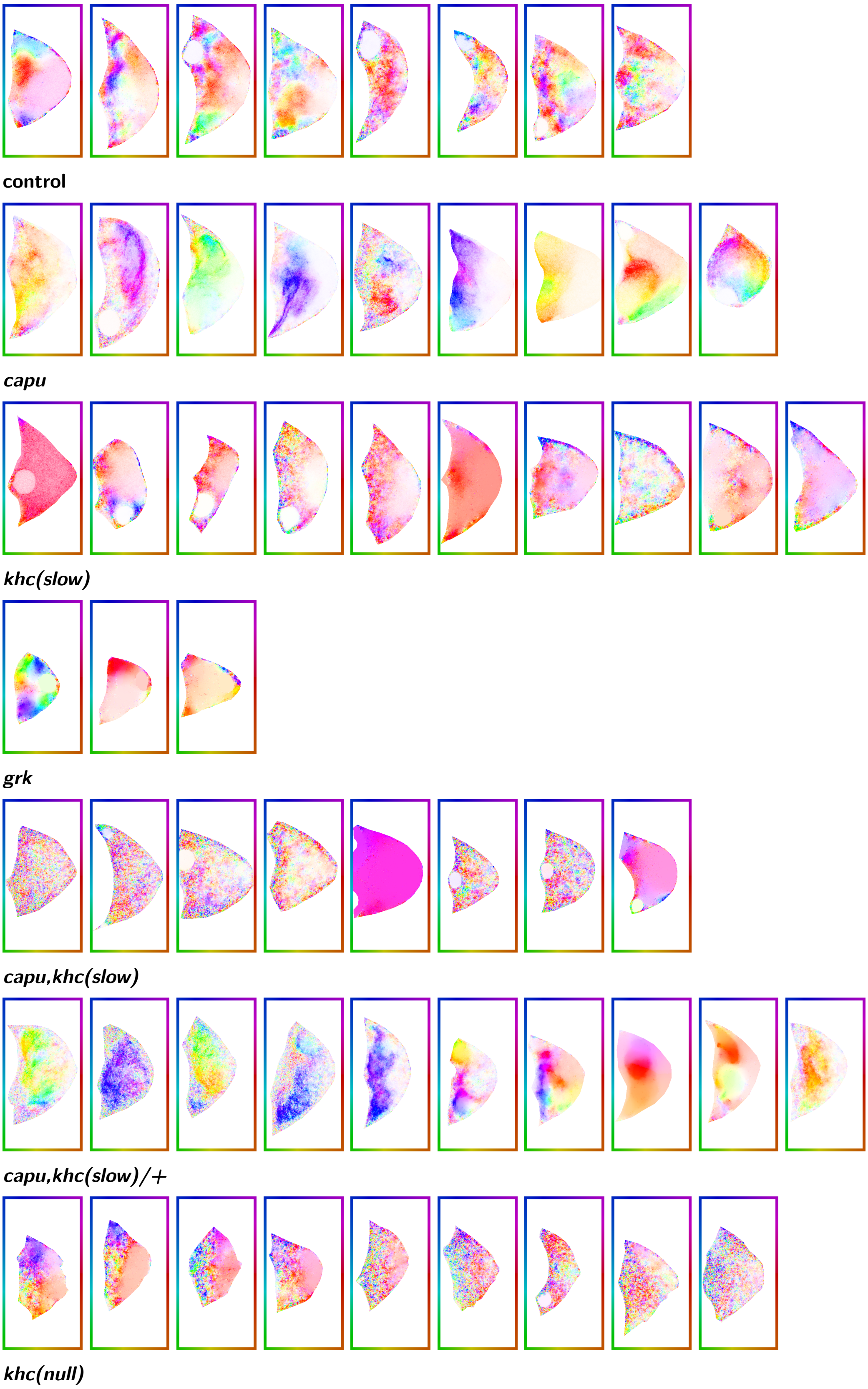


**Figure S5 - Mean velocity fields of oocytes in all analysed sequences.** The figure shows the estimated mean velocities (over time) for each sequence within the corresponding segmentation of the oocyte. All results were obtained with the parameter setting indicated in Suppl. Figs. S2 and S3.

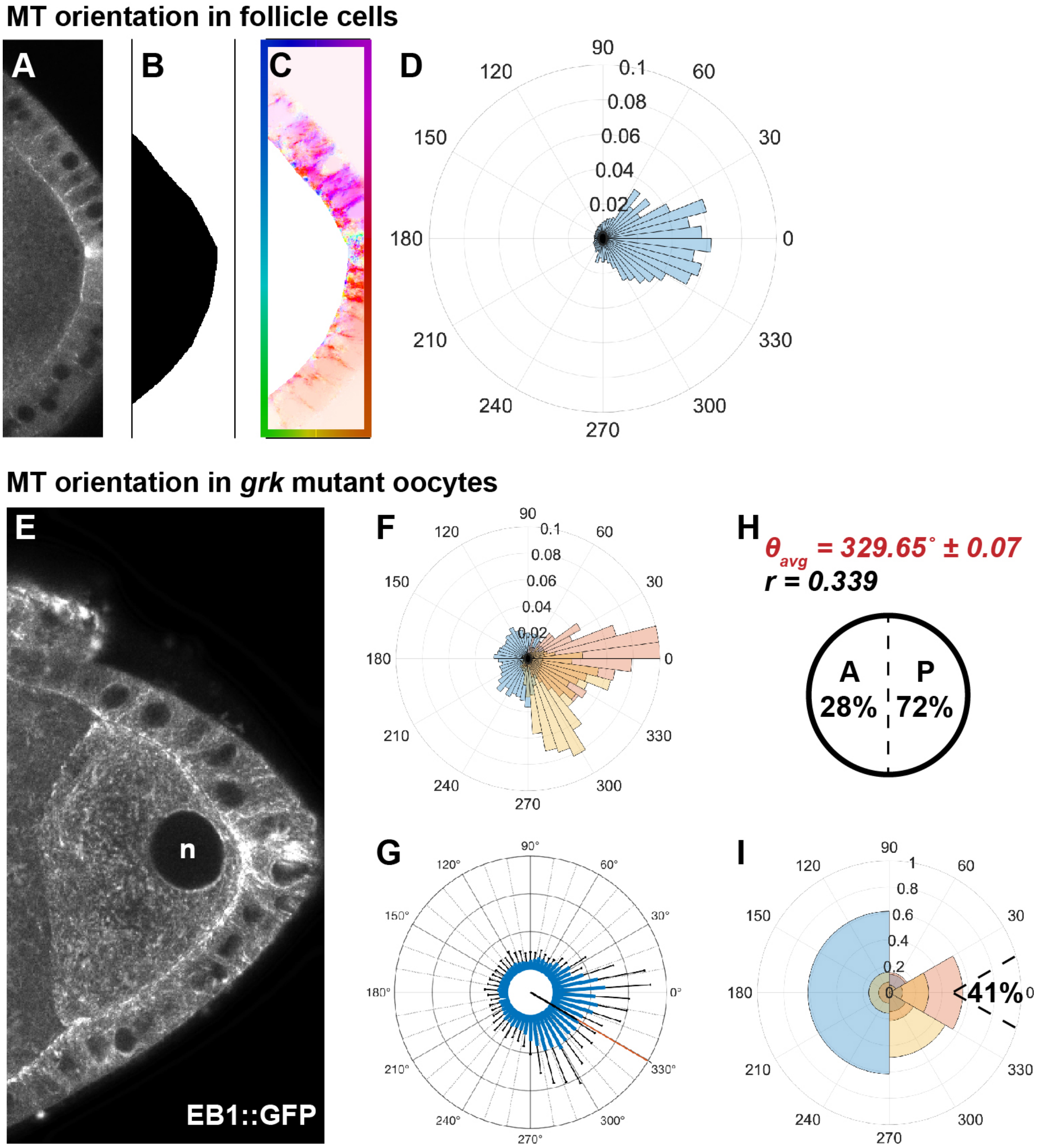


**Figure S6 - Validation of OF-based motion estimation of EB1 comet directionality and *grk* mutant oocytes. A)** First frame of the denoised sequence $u$. **B)** Hand-drawn segmentation mask of the oocyte (in black). **C)** Mean velocity (over time) outside the segmented region. **D)** Angular histogram based on the directions $\theta$ of the velocities $\mathbf{v}$ outside the oocyte. The shown result was obtained with parameters set to $\alpha_1 = 0.005$, $\beta_1 = 0.75$, $\alpha_2 = 0.0005$, and $\beta_2 = 0.005$. **E)** EB1 comets in *grk* mutant oocytes (standard deviation projection over 100 frames). Note the detached nucleus (n). **F)** Angular histograms of all analysed *grk* mutant oocytes. The distribution of angles of each analysed cell is shown in a different colour. **G)** Angular histogram of aggregated angles of all *grk* cells. Error bars indicate standard deviation for each bin. The red radial line indicates the mean angular direction $\theta_{\text{avg}}$ (average angle) and the black radial line the mean resultant vector; see (S13). **H)** Average angle of EB1 directionality in all *grk* mutant oocytes and the length $r$ of the mean resultant vector (indirectly proportional to the variance of the distribution). Moreover, the average posterior bias (72%) of all *grk* cells is shown. **I)** Angular histogram with only four bins ($30°$-$90°$, $90°$-$270°$, $270°$-$330°$, $330°$-$30°$) and average posterior tip directionality (41%).

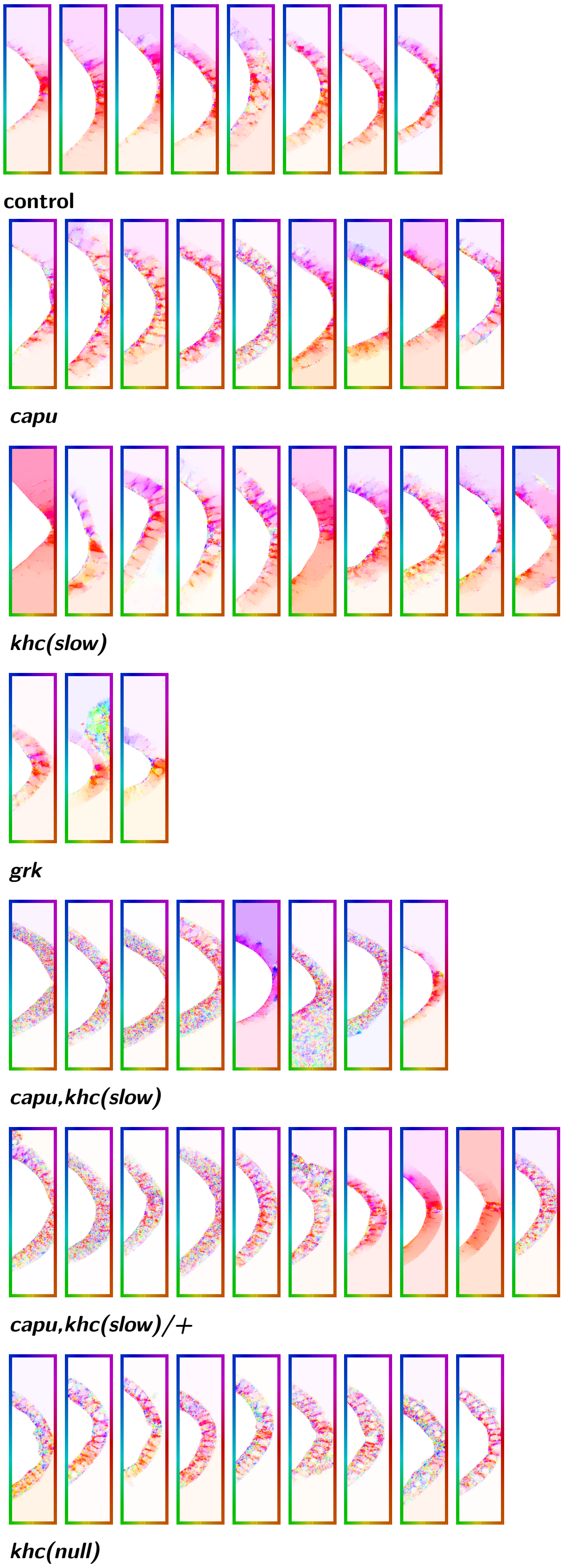


**Figure S7 - Mean velocity fields of follicle cells in all analysed sequences.** The figure shows the mean velocity (over time) in the follicle cells region for all datasets. All results were obtained with the parameter setting indicated in Suppl. Figs. S2 and S3.

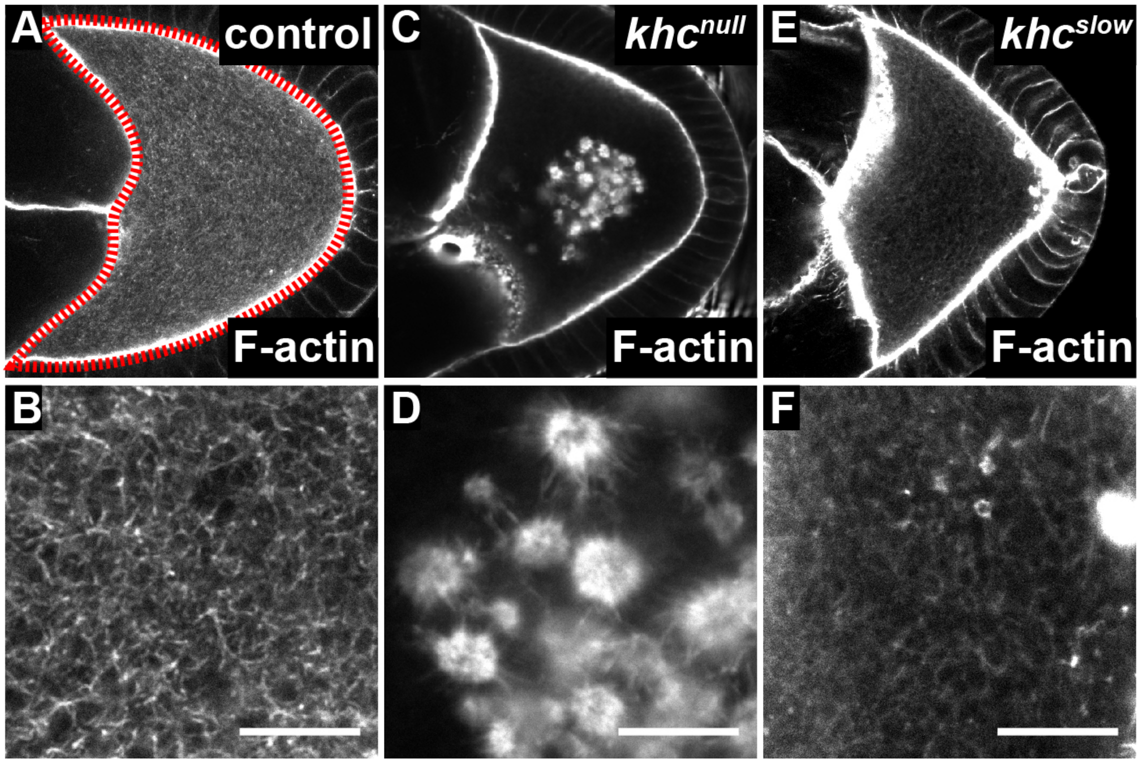


**Figure S8 - Actin mesh morphology in control and Kin mutant oocytes. A,B)** F-actin in fixed oocyte (encircled in red) traverses the entire cytoplasm. **C,D)** Oocyte, mutant for *khc*$^{27}$*(null)*, displays a substantial reduction of cytoplasmic actin filaments but exhibits large F-actin bearing vesicles. **E,F)** In comparison, *khc*$^{23}$*(slow)* mutant oocytes show a normal actin mesh morphology. Scale bars represent $10\,\mu\text{m}$.

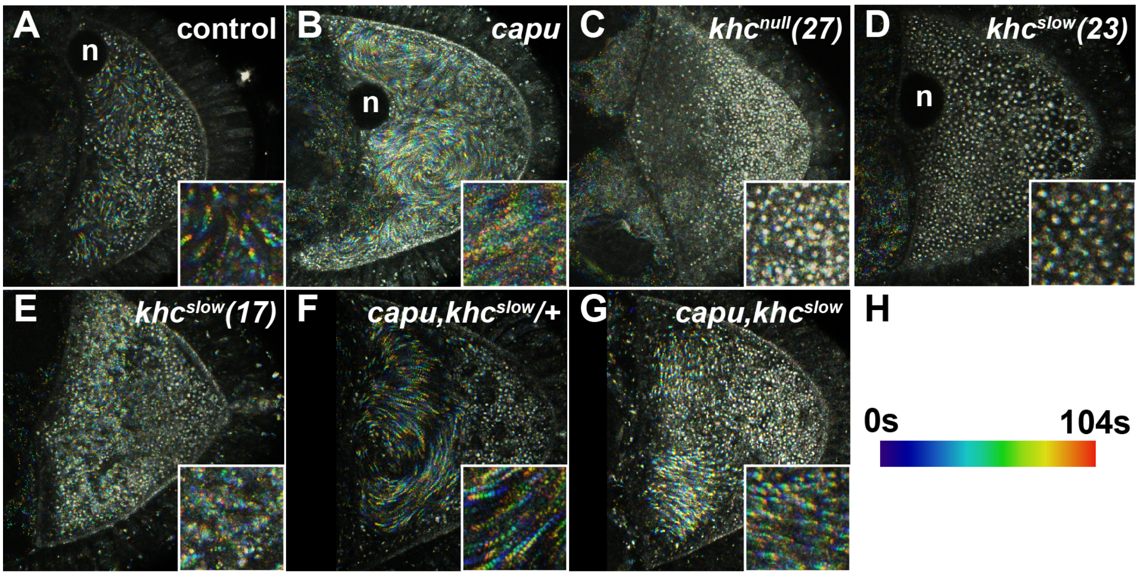


**Figure S9 - Cytoplasmic flows in control and mutant oocytes. A-G**) Stage 9 oocytes of control (A), *capu*$^{EY12344}$ (B), *khc*$^{27}$ (C), *khc*$^{23}$ (D), *khc*$^{17}$ (E), *capu*$^{EY12344}$/*capu*$^{EY12344}$,*khc*$^{17}$/+ (F) and *capu*$^{EY12344}$/*capu*$^{EY12344}$,*khc*$^{17}$/*khc*$^{17}$ (G) mutant oocytes. Auto-fluorescent vesicles were imaged as tracers to reveal cytoplasmic motion within the cells. Each image shows maximum intensity projections (colour coding in H) of ten consecutive frames. The insets depict a $15 \times 15\,\mu\mathrm{m}^2$ area of each cell to highlight presence or absence of cytoplasmic flows. If visible, nuclei are labelled (n).

#### Supplementary Movies

**Movie S1 - MT bulk motion in control and *capu* mutant oocytes (related to Fig. 1).** Dynamic bulk motion of Jup-labelled MTs in control (left) and $capu^{EY12344}$ mutant oocytes. Images have been acquired at one frame every $10.4\,\mathrm{s}$. Playback speed is $20\,\mathrm{fps}$ ($\sim 200\times$ real speed).

**Movie S2 - Example of image denoising of EB1 dynamics.** Confocal image sequence of dynamic EB1 comets before (left) and after (right) image denoising. Images have been acquired at one frame every $0.65\,\mathrm{s}$. Playback speed is $40\,\mathrm{fps}$ ($\sim 20\times$ real speed).

**Movie S3 - OF motion estimation in follicle cells.** Confocal image sequence of dynamic EB1 comets in follicle cells and subsequent OF motion estimation. Images have been acquired at one frame every $0.65\,\mathrm{s}$. Playback speed is $40\,\mathrm{fps}$ ($\sim 20\times$ real speed).

**Movie S4 - Dynamic behaviour of EB1 comets in *grk* mutant oocytes.** Denoised confocal image sequence of dynamic EB1 comets in a $grk^{2B6}/grk^{2E12}$ mutant oocyte. The nucleus (n) of the cell is detached and fails to migrate to the antero-lateral cortex. Images have been acquired at one frame every $0.65\,\mathrm{s}$. Playback speed is $40\,\mathrm{fps}$ ($\sim 20\times$ real speed).

**Movie S5 - Dynamic behaviour of EB1 comets in control and *capu* mutant oocytes.** Denoised confocal image sequence of dynamic EB1 comets in control (left) and $capu^{EY12344}$ mutant oocytes (right). Images have been acquired at one frame every $0.65\,\mathrm{s}$. Playback speed is $40\,\mathrm{fps}$ ($\sim 20\times$ real speed).

**Movie S6 - MT bulk motion in control and different Kin mutant oocytes.** Dynamic bulk motion of Jup-labelled MTs in control (first from left), $khc^{27}$ (second from left), $khc^{23}$ (third from left), and $khc^{17}$ (right) mutant oocyte. Images have been acquired at one frame every $10.4\,s$. Playback speed is $20\,\mathrm{fps}$ ($\sim 200\times$ real speed).

**Movie S7 - Dynamic behaviour of EB1 comets in control and different Kin mutant oocytes.** Denoised confocal time series of dynamic EB1 comets in control (left), $khc^{23}$ (middle), and $khc^{27}$ (right) mutant oocytes. Images have been acquired at one frame every $0.65\,\mathrm{s}$. Playback speed is $40\,\mathrm{fps}$ ($\sim 20\times$ real speed).

**Movie S8 - MT bulk motion in $capu,khc^{slow}/+$ and $capu,khc^{slow}$ double mutant oocytes.** Dynamic bulk motion of Jup-labelled MTs in $capu^{EY12344},khc^{17}/capu^{RK}$,GFP (left, transheterozygous mutant for $capu^{EY12344}/capu^{RK}$ and heterozygous for $khc^{17}$) and $capu^{EY12344},khc^{17}/capu^{RT}$, $khc^{17}$ (right, transheterozygous mutant for $capu^{EY12344}/capu^{RK}$ and homozygous for $khc^{17}$). Images have been acquired at one frame every $10.4\,\mathrm{s}$. Playback speed is $20\,\mathrm{fps}$ ($\sim 200\times$ real speed).

**Movie S9 - Dynamic behaviour of EB1 comets in $capu,khc^{slow}/+$ and $capu,khc^{slow}$ double mutant oocytes.** Denoised confocal time series of dynamic EB1 comets in $capu^{EY12344},khc^{17}/capu^{RK}$,GFP (left, transheterozygous mutant for $capu^{EY12344}/capu^{RK}$ and heterozygous for $khc^{17}$) and $capu^{EY12344},khc^{17}/capu^{RT}$, $khc^{17}$ (right, transheterozygous mutant for $capu^{EY12344}/capu^{RK}$ and homozygous for $khc^{17}$). Images have been acquired at one frame every $0.65\,\mathrm{s}$. Playback speed is $40\,\mathrm{fps}$ ($\sim 20\times$ real speed).

**Movie S10 - Recruitment of Kin towards MTs in the absence of the actin mesh.** A truncated KHC fusion protein (aa 1-700, C-terminal GFP fusion), localised in a posterior cloud in control oocytes (left), and was detected to weakly associate with filamentous MTs in the anterior region of the oocyte. Loss of *capu* caused a re-localisation of the protein, which is could now be found to heavily decorate MTs in the entire oocyte (right). Images have been acquired at one frame every $10.4\,\mathrm{s}$. Playback speed is $20\,\mathrm{fps}$ ($\sim 200\times$ real speed).
